# The conserved Pelado/ZSWIM8 protein regulates actin dynamics by promoting linear actin filament polymerization

**DOI:** 10.1101/2022.02.05.479255

**Authors:** Claudia Molina Pelayo, Patricio Olguin, Marek Mlodzik, Alvaro Glavic

## Abstract

Actin filament polymerization can be branched or linear, which depends on the associated regulatory proteins. Competition for Actin monomers occurs between proteins that induce branched or linear actin polymerization. Cell specialization requires the regulation of actin filaments to allow the formation of cell-type specific structures, like cuticular hairs in *Drosophila*, formed by linear actin filaments. Here, we report the functional analysis of CG34401/*pelado*, a gene encoding a SWIM domain containing protein, conserved throughout the animal kingdom. Mutant *pelado* epithelial cells display actin hair elongation defects. This phenotype is reversed by increasing Actin monomer levels or by either pushing linear actin polymerization or reducing branched actin polymerization. The same behavior occurs in hemocytes, where Pelado is essential to induce filopodia, a linear actin-based structure. We further show that this function of Pelado, ZSWIM8 in mammals, is conserved in human cells, where Pelado inhibits branched actin polymerization in a cell migration context. In summary, our data indicate that the function of Pelado/ZSWIM8 in regulating actin cytoskeletal dynamics is conserved, favoring linear actin polymerization at the expense of branched filaments.

## Introduction

The regulation of actin dynamics is essential in each cell to establish cell shape and drive cell division, migration, or morphological changes of whole tissues. Actin polymerization is regulated by several proteins that bind to either monomeric or filamentous Actin (Bogdan et al., 2013; Campellone and Welch, 2010; Carlier and Shekhar, 2017; Davidson and Wood, 2016; Hohmann and Dehghani, 2019; Revenu et al., 2004; Rottner et al., 2017; Vitriol et al., 2015). Over 150 proteins have been described to date, to participate as Actin binding proteins, promoting, or inhibiting specific features of its dynamic polymerization (Bogdan et al., 2013; Chen et al., 2014; Fletcher and Mullins, 2010; Frankel and Moosekert, 1994; Fricke et al., 2009; Goley and Welch, 2006; Gurung et al., 2016; Ito et al., 2011; Kovar, 2006; Luo et al., 1997; Machesky et al., 1999; Mullins et al., 1998; Pollitt and Insall, 2009a; Rotty et al., 2012; Takenawa and Suetsugu, 2007; Zallen et al., 2002). Actin polymerization can be classified as either branched or linear, depending on the associated regulatory proteins (Campellone and Welch, 2010; Suarez and Kovar, 2016). Linear actin polymerization generally requires the activity of the Formin protein family, like Diaphanous/Dia (Bogdan et al., 2013; Campellone and Welch, 2010), while branched polymerization requires the activation of the Arp2/3 complex, which is regulated among others, by Nucleator Promoting Factors (NPFs) from the WASP family, including Scar/WAVE or Wasp (Campellone and Welch, 2010; Clainche and Carlier, 2008; Frankel and Moosekert, 1994; Gautreau et al., 2021; Kelber and Klemke, 2011; Mullins et al., 1998). Although cells have a high concentration of actin monomers (Burke et al., 2014; Koestler et al., 2009; Pollitt and Insall, 2009a), most of it is sequestered by these different regulatory proteins, establishing a competition for actin monomers between the two distinct forms of polymerization (Burke et al., 2014; Henson et al., 2015; Suarez and Kovar, 2016; Suarez et al., 2015). In most cells there is a larger number of proteins that induce branched actin polymerization, and thus competition for actin monomers is often tilted towards branched polymerization arrangements (Suarez et al., 2015). Profilin (*chickadee/chic* in *Drosophila*) is pivotal in the polymerization selection process with Chic/Profilin both favoring linear actin polymerization, mediated by the Formin protein family, and also inhibiting branched polymerization by preventing the binding of NPFs to Actin monomers (Suarez et al., 2015).

Cell specialization requires actin cytoskeleton regulation to allow the formation of specific structures, like for example cuticular hairs in *Drosophila*. Cuticular epithelial cells in *Drosophila* form a single cuticular actin hair at the apico-distal vertex of each cell. Hair formation depends on the Frizzled Planar Cell Polarity (Fz/PCP) pathway, which controls the positioning and polarized Actin accumulation by regulating the hierarchical activity of groups of genes that constitute it: core PCP genes (*frizzled/fz, disheveled/dsh, diego/dgo, Van gogh/Vang, prickle/pk*, and *flamingo/fmi*), and Planar Polarity Effector (PPE) genes, which either inhibit actin polymerization at the proximal side of wing epithelial cells (e.g. *inturned, fritz, fuzzy*, and *multiple wing hairs*/*mwh*), or promote actin polymerization as positive effectors of the Fz/Dsh/Dgo complex (Adler, 2012; Carvajal-Gonzalez and Mlodzik, 2014; Gao, 2012; Strutt and Strutt, 2009; Winter et al., 2001; Yan et al., 2009). One of these downstream effectors of the Fz/Dsh complex is RhoA (a.k.a. Rho1 in *Drosophila*), which locally activates its effector proteins, including in this context the *Drosophila* Rho-associated kinase, Drok. Drok activity focuses actin polymerization, but does not affect the location or orientation of wing hairs (Adler, 2012; Carvajal-Gonzalez and Mlodzik, 2014; Gao, 2012; Strutt and Strutt, 2009; Winter et al., 2001; Yan et al., 2009). Actin hair localization and formation is widely used as a proxy to evaluate the Fz/PCP pathway functionality (Adler, 2012; Carvajal-Gonzalez and Mlodzik, 2014; Carvajal-Gonzalez et al., 2016; Gao, 2012; Gault et al., 2012; Lee and Adler, 2004; Lu et al., 2010; Mitchell et al., 1983; Sagner et al., 2012; Strutt and Strutt, 2009).

Actin hair formation also requires the activity of the Shavenbaby/Shavenoid pathway that regulates the actin cytoskeleton (Delon et al., 2003; Ren et al., 2006). Since the hair structure, trichome, is formed by linear actin polymerization, it is a useful and well characterized model system to identify and analyze proteins that regulate actin dynamics (Adler, 2018; Adler et al., 2013; Eaton et al., 1996; Guild et al., 2005; He et al., 2005; Matis et al., 2014; Ren et al., 2007; Turner and Adler, 1998; Winter et al., 2001; Yan et al., 2009). In particular, the wing epithelial cells in *Drosophila* are uniquely suited for such analyses, as all wing cells form a single actin hair (or trichome) at the distal apical vertex of each cell (Adler, 2012; Klein et al., 2006; Mitchell et al., 1983; Sagner et al., 2012; Singh and Mlodzik, 2012; Strutt and Strutt, 2009). As this structure is highly repetitive and requires a specific and exquisite interplay of regulatory factors of actin dynamics, it serves as an excellent model to identify and investigate novel genes required in this process (Adler, 2018; Adler et al., 2013; Eaton et al., 1996; Fagan et al., 2014; Fang and Adler, 2010; Guild et al., 2005; He et al., 2005; Ren et al., 2007; Turner and Adler, 1998). Importantly, this process is critically linked to linear actin polymerization and the associated formin, Diaphanous (Dia) (Mitchell et al., 1983; Lu and Adler, 2015). Despite accumulating knowledge of the mechanisms and proteins regulating polarized actin accumulation during trichome formation, many open questions remain and particularly the mechanisms regulating linear versus branched actin polymerization in the cuticular epithelial cells, and in other contexts, remains largely unknown (Delon et al., 2003; Fang and Adler, 2010; Fang et al., 2010; Gault et al., 2012; Guild et al., 2005; He et al., 2005; Ren et al., 2005, 2007; Turner and Adler, 1998).

Here, we describe the identification, and functional characterization of CG34401/*pelado* (*pldo*), a gene encoding a multidomain protein that is conserved throughout the animal kingdom, called ZSWIM8 in mammals (Shi et al., 2020). We show that it is required for the formation of the actin-based trichome formation in *Drosophila* cuticle cells, most notably on wings and the notum. This gene has also been characterized in *C. elegans*, as an E3 ubiquitin ligase, named EBAX-1, and is part of a ubiquitin ligase complex that includes Elongins B and C, and Cullin2 (Wang et al., 2013). Although this E3 ligase function was also described in specific cell lines, the gene being named *dorado* in this case (Shi et al., 2020), our data suggest that in the context of actin dynamics Pldo/ZSWIM8 may act independently of its E3 ligase activity.

Our functional genetics and epistasis analyses demonstrate that Pldo favors linear actin polymerization – at the expense of branched polymerization - in *Drosophila* and human cell lines, and thus that this function of Pldo in regulating the actin dynamics is conserved. Immunoprecipitation assays suggest that Pldo is part of a multiprotein complex that includes Scar/WAVE, Diaphanous, and Profilin (Campellone and Welch, 2010; Hohmann and Dehghani, 2019a; Rottner et al., 2017; Suarez and Kovar, 2016). Structure function studies reveal that the N-terminal region of Pldo, containing the highly conserved SWIM, FH1 and WH2 domains, is sufficient to promote the formation/induction of filopodia in general, and that the C-terminal part might be required for cell type specific actin related functions, for example in epithelial cells. Noteworthy, the *pelado* gene was also identified in a screen related to human diseases (microcephaly) (Yamamoto et al., 2014), without a described mechanistic function in that context, which makes it a very interesting protein to analyze. Consistent with our data on actin regulation established below, *pldo* function in the central nervous system might be related to axon/dendrites formation, or radial glial migration that give rise to the neurons that populate the different brain layers, being all processes that requires specific actin polymerization regulation (Chou and Wang, 2016; Cooper, 2013; Kullmann et al., 2020; Rust et al., 2012). Taken together, our studies suggest that *pldo* is an essential regulator of actin dynamics in *Drosophila* and human cells, with a likely link in the disease context and even potential roles related to neurodegenerative diseases.

## Results

### Pldo, a novel gene required for cellular hair formation in *Drosophila*

In an ENU mutant suppressor genetic screen, designed to identify the gene responsible for a wing growth phenotype produced by the overexpression drive by an EP line (EP361, (Cruz et al., 2009)), we generated a recessive embryonic lethal mutation on the X chromosome. To study the cellular phenotype in homozygous mutant cells, we used the Mosaic Analysis with a Repressible Cell Marker (MARCM) technique (Wu and Luo, 2006). Strikingly, homozygous mutant epithelial cells from different adult tissues, including the wing and notum, displayed a lack of trichome (cellular actin-based hairs) phenotype (Figure 1A-C; also Suppl. Figure S1A-B). To identify the affected locus, we employed first a complementation strategy using several genomic duplications spanning the X chromosome, followed by small deficiencies to further restrict the chromosomal region associated with the phenotype. Subsequently, we knocked down the expression of candidate genes present within this chromosomal region by using dsRNA constructs and screened for the absence of trichomes, as seen in the phenotype obtained in homozygous mutant wing cells (Suppl. Figure S1D and H). This approach identified the previously uncharacterized gene CG34401 (specific target region of the RNAi construct is shown in Figure 1D) as responsible for the “loss of hairs” phenotype. As the defect associated displayed a hairless, or “bald”, phenotype, we named this gene *pelado* (*pldo*, which is the Spanish word for “bald”).

**Figure 1:**
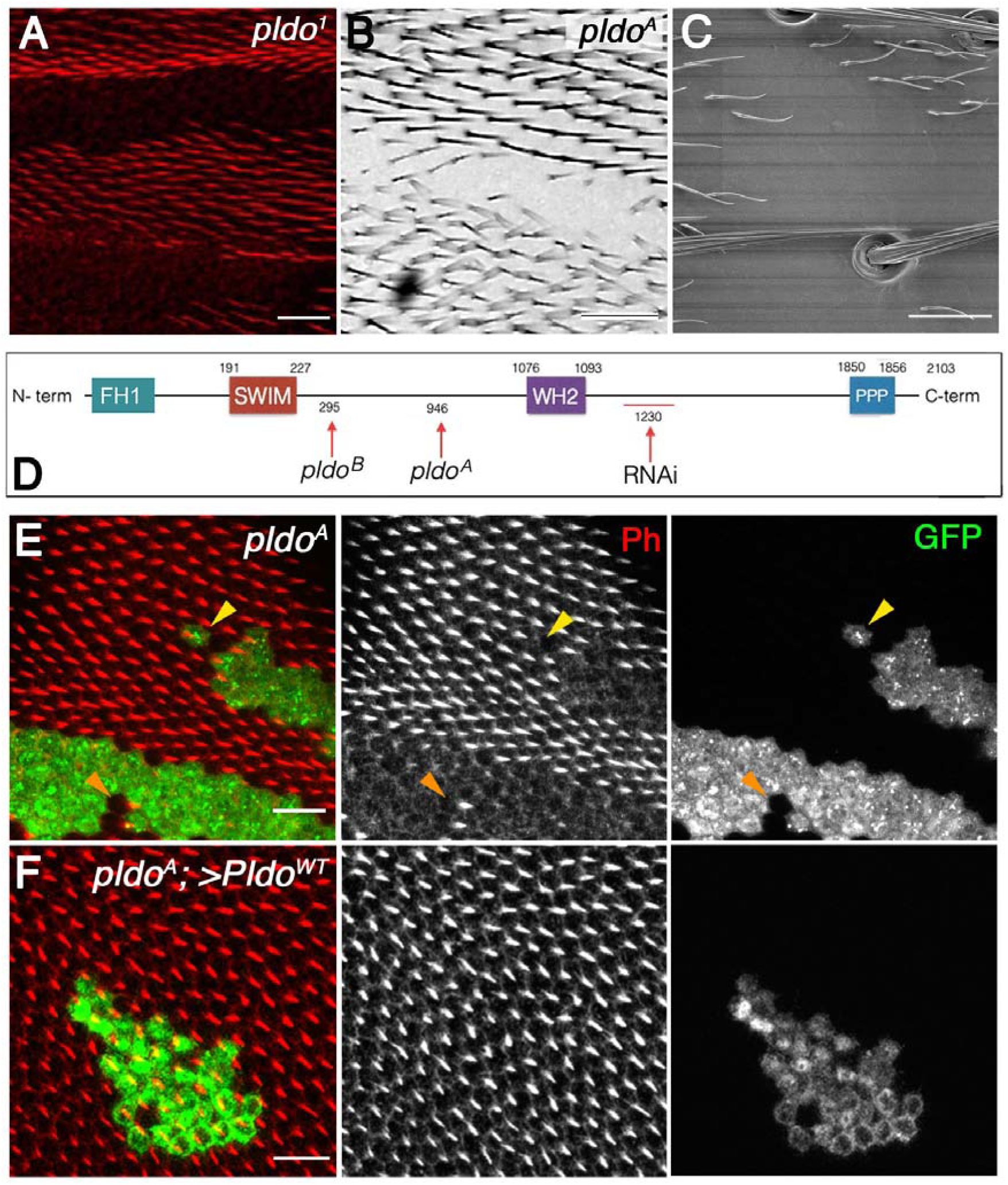
Pldo is required for epithelial cuticle hair formation. A-D. General description of CG34401/*pldo* mutant phenotypes and protein sequence. Unmarked *pldo*^*1*^ clones showing the absence of trichomes (actin hairs) in a pupal wing (A), adult wing (B, *pldo*^*A*^ clone), and adult thorax (C, SEM micrograph of thorax with *pldo*^*1*^ clone). Scale bars correspond to 100μm in A, and 50μm in B and C. Note that cuticular cellular hairs are missing, but that sensory bristles are not affected. Schematic presentation of Pldo protein sequence (D) with the main conserved domains indicated (residue numbers are above the protein sequence line). Below the line are the characterized mutant alleles indicated, *pldo*^A^and *pldo*^B^ (available in the Bloomington Stock Center), all showing the same phenotypes. These two alleles have point mutations: *pldo*^A^: G18874432A generates a non-sense mutation (W946Stop codon); and *pldo*^B^: C18872412T generating the change T295M. The RNAi construct sequence (BL 18553) is also shown. E. Pupal wings with MARCM *pldo* mutant clones (mutant cells marked in green with GFP). Cellular hairs were visualized by rhodamine-phalloidin staining (red). Note that the *pldo* LOF phenotype is fully cell autonomous, as observed in either a single cell clone (yellow arrowhead) or a two-cell wildtype patch (orange arrowhead) displaying the absence or presence of hairs, respectively. Scale bar correspond to 50 μm. F. *pldo* loss-of-function hair phenotype is fully rescued by the expression of Pldo^WT^ (via UAS in the MARCM clones), with hairs showing normal polarity and length. Scale bar represents 50 μm.

The CG34401/*pelado* gene (*pldo*) is located on the X chromosome at 17F2 and encodes a large protein of 2103 amino acids and approx. 250kDa in size (Figure 1D). *In silico* analysis of the Pldo sequence (see Materials and Methods) revealed the presence of several conserved domains (Figure 1D). The SWIM domain (Swi2/SNF2 and MuDR zinc-binding), which is defined as a domain that can interact with both DNA and other proteins (Godin et al., 2015; Ko et al., 2014), gives the name to the human ortholog of Pldo, ZSWIM8 (Zinc finger SWIM-type containing 8; www.flybase.org). Interestingly, other conserved domains present in Pldo include the FH1 (Formin Homology 1 domain), a Wiskott-Aldrich homology 2 domain (WH2), and a Polyproline region (Figure 1D, also Suppl. Figure S1K), which are also present in several proteins that regulate actin cytoskeleton dynamics, including Scar/WAVE and the formin Diaphanous/Dia (Carlier et al., 2013; Miki et al., 1998; Paunola et al., 2002; Watanabe et al., 1997). In particular, the WH2 domain binds to monomeric Actin (Paunola et al., 2002), and FH1 and Polyproline domains can bind to Profilin (Chicadee/Chic in *Drosophila*) (Faix and Grosse, 2006).

Following the identification of *pldo* as the gene of interest, we identified two additional mutant alleles via complementation analyses, indicating that those mutations are affecting the same gene. These alleles were generated in a genetic screen in the lab of Hugo Bellen in a screen for mutants related to human diseases (Haelterman et al., 2014; Yamamoto et al., 2014); available in the Bloomington Drosophila Stock Center/BDSC with stock numbers 52333 and 52334. We subsequently refer to these molecularly characterized alleles as *pldo*^*A*^ and *pldo*^*B*^, respectively (Figure 1D), with one being a non-sense mutation that produces a stop codon at residue 946, W946STOP (*pldo*^*A*^), and the other a point mutation, T295M (*pldo*^*B*^; see Figure 1D for details). In accordance with the analysis of the original allele identified in our laboratories (*pldo*^*1*^), homozygous mutant clones of these alleles also displayed loss of cellular hairs/trichomes (Figure 1E and Supplementary Figure S1A-B). This phenotype is cell autonomous (see examples in Figure 1E, with very small clones displaying the respective features). To unequivocally confirm the identity of the gene responsible for the loss of trichomes, we generated a transgenic line to express the *pldo*^*WT*^ coding sequence in the *pldo*^A^ mutant background using the MARCM technique (Wu and Luo, 2006). Indeed, the expression of the Pldo^WT^ transgene completely rescued the hair loss defects, with no other evident phenotype in the mutant clones (Figure 1F).

As *pldo* is a recessive lethal mutation, we next wished to establish the stage of lethality. To determine this, we generated *pldo* mutant stocks balanced with FM7 marked with GFP (expressed under the *twist* driver), and thus GFP negative embryos or larvae would be homozygous mutant for *pldo*. The number of GFP-negative embryos (*pldo* mutants) was considerably lower than controls. We were also able to observe a small number of mutant larvae (at 24 hours AEL [after egg-laying]) with more than half of these being already dead, and the ones still alive showed severe motor defects (see summary of these results in Suppl. Figure S1I-J). Taken together with maternal expression of *pldo* (www.flybase.org), these results suggest an important function of *pldo* during early embryonic developmental stages, which is obscured by maternal contribution.

### Pldo is required for actin hair formation and elongation

Actin hair formation in cuticular epithelial cells takes place in three stereotyped stages: Apical actin accumulation, prehair formation, and hair elongation (Guild et al., 2005). Each stage is characterized by specific Actin distributions and hair features. At around 28 hours After Puparium Formation (APF), Actin starts accumulating apically in the distal vertex of every epithelial wing cell (Mitchell et al., 1983; Turner and Adler, 1998). At 32 hours APF, the structure called “prehair” appears, and finally at 36 hours APF, the hair is almost completely formed or elongated (Classen et al., 2008; all time points relate to a growing temperature of 25ºC). To address the potential role of Pldo in these stages, we generated *pldo* mutant clones at third instar larvae and followed the cellular maturation and hair growth in the wing epithelia at the time points of pupal development mentioned above. While at 28 hours APF, there were no obvious difference in Actin distribution between *pldo* clones and control cells (Figure 2A), there were clear alterations in Actin distribution and (pre)hair formation at 32 and 36 hours APF (Figure 2B-C). In *pldo* mutant cells, the prehair formation process was impaired, and defects were most evident from early stages of hair elongation. Thus, the core PCP-regulated hair formation process, which requires localized linear actin polymerization (Lu and Adler, 2015; Mitchell et al., 1983), was severely impaired in the *pldo* mutant cells.

**Figure 2:**
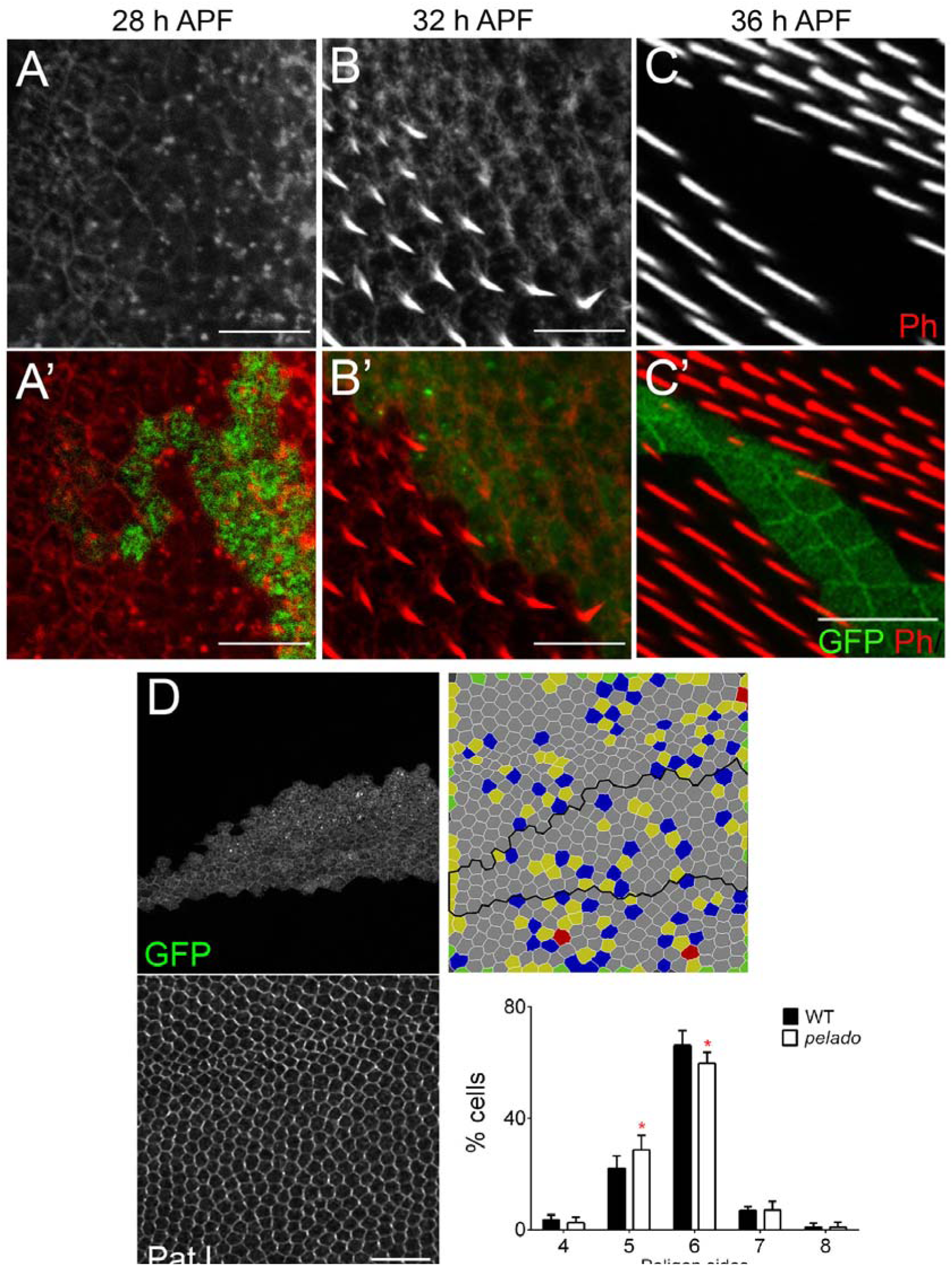
Pldo is required for actin hair formation and cell shape maturation. A-C. Temporal characterization of the mutant *pldo* phenotype during cellular hair formation. Confocal micrographs showing *pldo* MARCM clones (marked by GFP, green) during actin hair formation (labelled via rhodamine-phalloidin). (A-C) monochrome images showing Actin alone, and (A’-C’) displaying clonal marker (green) and Actin (red). Note absence of hair elongation from 30-32hrs APF onward, in mutant GFP marked cells. (A-A’) Note accumulation of Actin appears normal in control and mutant cells at 28h APF. All the images are oriented with distal in the lower right corner. Scale bars correspond to 50 μm. D. Evaluation of cell shape in *pldo* mutant cells, at 28-32 h APF, as compared to neighboring wild-type cells. Note the significant reduction in the number of hexagonal cells in *pldo* mutant clones (marked by GFP; cell outline displayed via PatJ staining, a junctional marker). 6 different individuals were analyzed, with around 100 cells in each case. *P*=0,0366 via t-test. Scale bar correspond to 100 μm.

To further address the role of *pldo* in the hair formation process, we tested its relationship to the PCP pathway, as Fz/PCP signaling is critical to focus actin polymerization to the distal vertex. Earlier stages of PCP establishment, preceding hair formation, are characterized by the asymmetric localization of its core components (e.g rev. in Adler et al., 2013; Carvajal-Gonzalez and Mlodzik, 2014; Matis et al., 2014; Strutt and Strutt, 2009). We thus evaluated if the asymmetric core PCP complex localization is affected. Fmi serves as the anchor of both distal and proximal PCP complexes, and its localization reflects this feature as a regular zig-zag pattern enriched in the proximal-distal axis, which can be visualized and quantified (Aigouy et al., 2010). We did not detect any changes in Fmi localization in wing cells mutant for *pldo*, as compared to neighboring control cells (Suppl. Figure 2), indicating that Pldo is neither upstream, nor part of the regulatory circuit of the core PCP pathway. This observation and interpretation is consistent with the phenotypes seen by altering the expression of core PCP factors or direct regulators of this pathway, with the actin hairs often being formed, but in abnormal number and/or locations (Lu et al., 2015; Yan et al., 2008).

A second feature regulated by the core PCP pathway, during pupal wing development and in general, is the organization of the epithelial cells themselves with their arrangement into a hexagonal pattern (Etournay et al., 2015; Sánchez-Gutiérrez et al., 2013; Wong and Adler, 1993). This terminal shape establishment follows a cell shape elongation along the proximo-distal axis, when PCP is reorganized from a radial polarity arrangement to a proximal-distal axis alignment (Aigouy et al., 2010). We thus asked whether *pldo* might be required in this context and compared the polygon distribution of *pldo* mutant cells to surrounding wild-type control cells in pupal wing cells at 28-32 hours APF. Interestingly, cells mutant for *pldo* display a reduction in the proportion of hexagonal shape, compared with surrounding control cells (Figure 2D), suggesting a role for Pldo in the cortical actin bundle function.

Taken together with the actin hair phenotype, these data suggest that Pldo likely acts as an effector of the core PCP pathway to regulate actin cytoskeleton associated functions like cell shape definition and actin accumulation during hair formation and elongation. Consistent with this notion, *pldo* encodes protein domains associated with Actin binding and regulation of its polymerization, such as WH2, FH1 and Polyproline domains.

### Pldo promotes linear actin polymerization during hair outgrowth

To gain insight into the underlying regulatory mechanism(s) of how Pldo might function in actin filament dynamics, we asked whether the *pldo* mutant phenotype can be genetically modified by genes encoding proteins regulating linear and/or branched actin polymerization. Such genes fall into the category of Formins, including Dia or the anti-formin Mwh, or the WASP family, including Scar/WAVE and Wasp, among others (a list of actin regulators tested is shown in Suppl. Figure S3C). Specifically, for example, Dia and Enabled/Ena promote linear actin polymerization, while Scar and Wasp induce branched actin polymerization by means of activating the Arp2/3 complex (Campellone and Welch, 2010; Carlier et al., 2013; Clainche and Carlier, 2008; Goley and Welch, 2006; Kang et al., 2010; Machesky et al., 1999; Paunola et al., 2002; Rotty et al., 2012). Mwh, a functional anti-formin inhibits Dia function, preventing linear actin polymerization (Lu and Adler, 2015). To assay for potential interactions, we expressed *pldo-IR* under the control of *ptcGal4* leading to a reduction of trichomes within the *ptc* expression domain, mimicking the *pldo* loss-of-function (LOF) phenotype. In this context, a phenotypic rescue of *pldo* was detected when increasing Dia or Actin protein levels in the *ptcGal4>pldo-IR* background (Figure 3B-C and Suppl Figure S3A). Strikingly, this *pldo* knockdown phenotype was also “rescued” by reducing Arp2, Arp3, Scar or Mwh levels (Figure 3B-C and Suppl Figure S3A-B); noteworthy, a *mwh* knockdown reversed the *pldo* LOF phenotype and produced the well described “multiple wing hair” defects, characterized by the appearance of more than one short hair per cell (Suppl Figure S3A-B) (Lu et al., 2015; Yan et al., 2008). As mentioned above, Shavenoid/Sha is another component required for trichome formation. *sha* LOF alleles show either the absence of hairs or the appearance of more than one tiny hair; in consequence, it is described as an inducer of linear actin polymerization (Adler, 2018; Delon et al., 2003; Ren et al., 2006). We asked whether increasing Sha protein levels allowed the hair to form in a *ptcGal4>pldo-IR* background. Indeed, Sha co-oveexpression in this background restored hair formation (Figure 3B-C). Interestingly, Pldo knockdown phenotypes in the antenna laterals (Suppl Figure S1F) is similar to the phenotype observed in *shavenoid* mutant flies (He and Adler, 2002). Importantly, these phenotypic rescues were also observed when assayed in *pldo* MARCM clones and analyzed in pupal wings (see Figure 3D for examples). In summary, promoting linear actin polymerization by increasing the levels of Dia, Ena, Sha, and Profilin/Chic in the *ptcGal4>pldo-IR* background restored hair formation and rescued the phenotype induced by *pldo* knock-down. A similar “rescue” was observed by reducing Mwh levels, an inhibitor of Dia activity.

**Figure 3.**
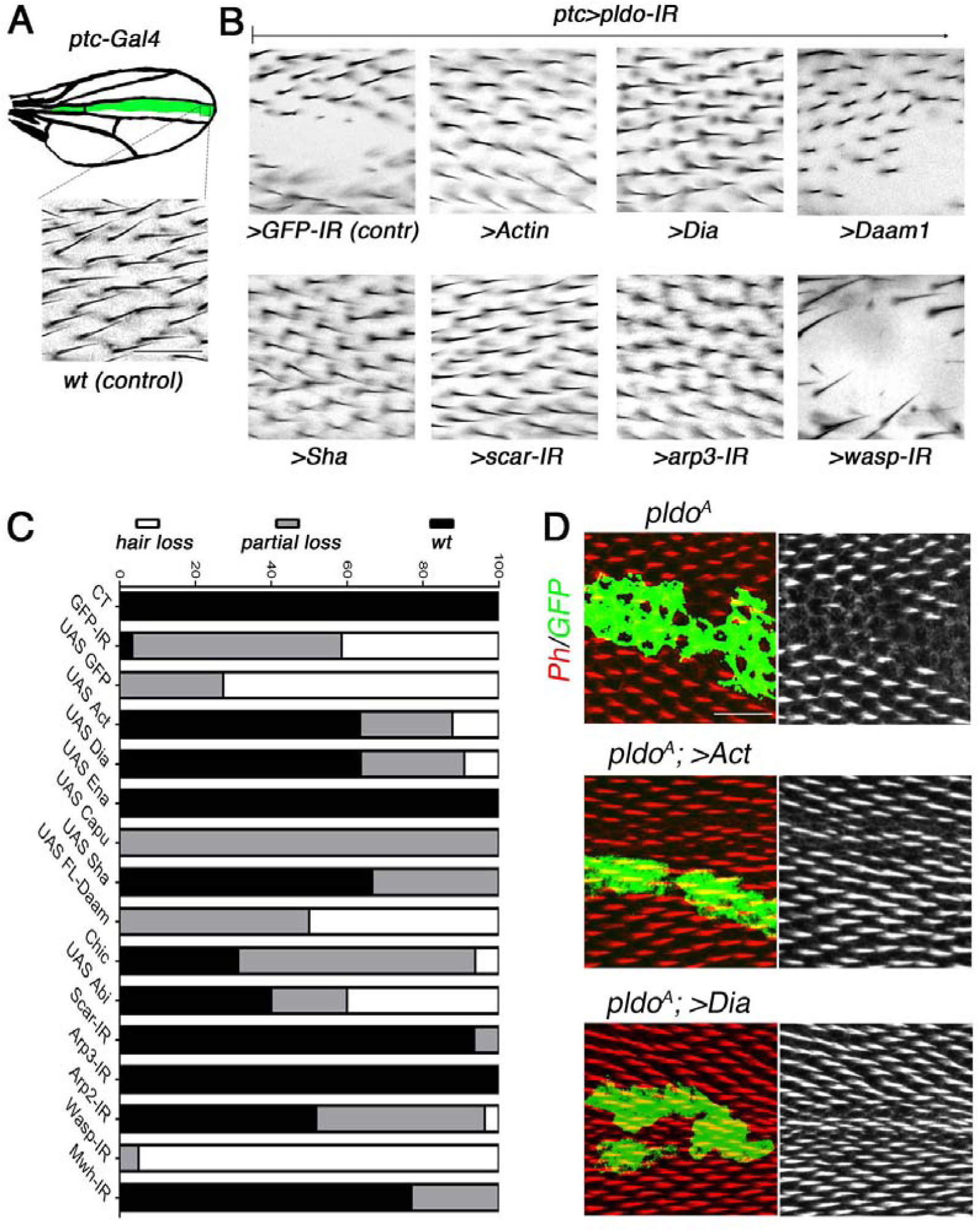
Pldo mediates competition for monomeric Actin during trichome formation. A. Schematic displaying the *ptc*Gal4 expression pattern (green area) used in pupal and adult wings. Square in drawing corresponds to region shown in lower panel (*wt* control) and (B) panels. Scale bar corresponds to 50 μm (also for panel B). B-C. Genetic interactions between *pldo-IR* (knock-down) and factors that regulate actin cytoskeleton polymerization with phenotypic appearance as evaluated in adult wings. Squares display high magnification views of wing area (boxed in schematic in A) expressing *ptcGal4, UAS-Dcr, UAS-pldo-IR* in combination with the indicated transgenes, either knocking-down additional factors or overexpressing a gene of interest. Compare control phenotype of *ptcGal4, UAS-Dcr, UAS-pldo-IR, UAS-GFP-IR* (upper left panel) to *wt* in (A) and the genotypes of interest (B). Potential reversion of the *pldo-IR* (LOF) phenotype was evaluated according to the size of the area in adult wings, which lost the trichomes. Quantification was performed by scoring each wing with either “wild-type” (wt, normal hair number), “partial loss of trichomes” (partial loss) or “complete loss of trichomes” (hair loss) as shown in graph in (C). Percentages of the respective appearance are shown in C, see (B) for phenotypic reference: complete reversion to wild-type as seen with *UAS-scar-IR*, and partial reversion displaying small areas with trichome loss as seen with *UAS-Capu* (Suppl Figure S3B), and no reversion (large area with no trichome) as seen in control *UAS-GFP-IR* or *UAS-wasp-IR*. Quantification is based on scoring >10 individuals in each case. D. Reversion of the *pldo* (LOF) phenotype as seen in *pldo* mutant MARCM clones in pupal wings to confirm the extent of phenotypic rescue. Note that Actin and Dia overexpression/gain-of-function (GOF) showed a complete reversion of *pldo* mutant clones in pupal wings. Scale bar corresponds to 25 μm.

Competition for Actin monomers in the regulation of linear versus branched actin polymerization has been described (Burke et al., 2014; Davidson and Wood, 2016; Davidson et al., 2018; Jaiswal et al., 2013; Suarez and Kovar, 2016; Suarez et al., 2015; Vitriol et al., 2015), and we thus asked whether a similar competitive nature exists during trichome formation. To address this, we asked whether reducing branched actin polymerization by knocking-down *scar, arp2, arp3*, and *wasp* has an effect in the *ptcGal4>pldo-IR* background. Strikingly, we observed a reversion of the phenotype in most cases, including *scar, arp2*, and *arp3* knockdowns (Figure 3B-C and Suppl. Figure S3A-B). The exception was *wasp* knock-down (Figure 3B-C). However, as Scar and Wasp have specific roles in different cell types (Berger et al., 2008; Koch et al., 2012; Rajan et al., 2009; Zallen et al., 2002), these results suggest that Scar function is relevant in pupal wing cells, but Wasp is not required there. Taken together, these results suggest that competition for Actin monomers during trichome formation in *Drosophila* is a critical component of its regulatory circuitry. To further corroborate this notion, we evaluated the effect of increasing Actin monomer levels in the *ptcGal4>pldo-IR* background. Strikingly, a *ptcGal4>pldo-IR, >Act* genotype displayed a complete reversion of the trichome loss (Figure 3B-C), consistent with the hypothesis that Pldo influences the competition for Actin monomers.

In summary, these data suggest that the function of Pldo is to promote linear actin polymerization required during trichome formation, which could be either through the inhibition of branched actin polymerization or by directly regulating linear actin plymerization. The combination of reduction or increase of Actin itself, or branched or linear actin filament promoting factors suggest that Pldo promotes linear actin polymerization by positively acting on its regulators, and/or represses branched actin regulators. As such, Pldo functions in this context as a balance tipping factor towards linear actin polymerization, either by a direct promotion of linear actin polymerization or by inhibiting branched actin polymerization, or both.

### Pelado is required for filopodia formation in hemocytes

To ask whether Pldo serves this function in general, we wished to evaluate its requirements in a different cell type. We thus asked whether *pldo* is required to regulate cell morphology in hemocytes, a migratory *Drosophila* blood cell similar to mammalian macrophages (Moreira et al., 2013) (Figure 4A). *pldo* mutant hemocytes (generated by the MARCM technique) displayed a reduced attachment cell area and a significant increase in the structures known as ruffles (Figure 4A, bottom panel, and Suppl Figure S4E). This phenotype was rescued by expressing Pldo^WT^ in this experimental setting (Figure 4B, rescued cell is marked by GFP). Exogenous expression of Pldo^WT^ in the *pldo* mutant cells, which is likely an overexpression relative to its endogenous levels, did in fact not just rescue the LOF defects, but caused an increase in the formation of filopodia-like protrusions (Figure 4B). This observation is consistent with its proposed role in favoring linear actin polymerization (which could be a direct effect or by preventing branched actin polymerization). Moreover, similar to the wing hair formation assay (Figure 3 above), the *pldo* hemocyte defects were also rescued by co-(over)expression of Dia, Actin, or by IR-mediated knock-down of *scar* (Suppl. Figure 4A-B).

**Figure 4:**
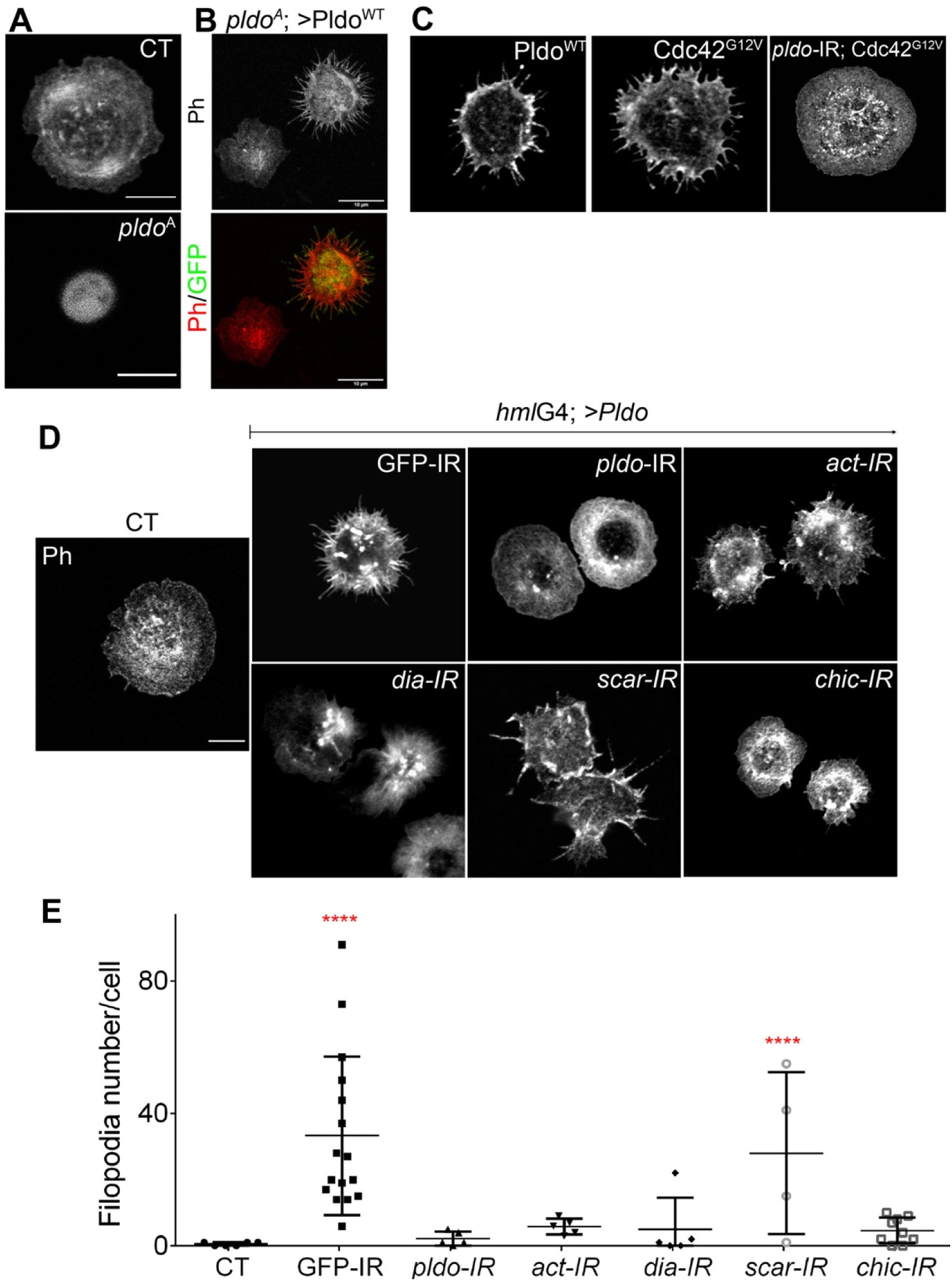
Pldo induces formation of filopodia in hemocytes. A. Confocal microscopy images of hemocytes stained with rhodamine-phalloidin to visualize actin filaments. Control wild-type hemocyte (CT, top) is shown for comparison to *pldo* mutant cells, which show a reduction in cell attachment area (*pldo*^A^, bottom panel). Scale bar represents 5 μm. B. Reduction in cell attachment area was rescued by co-expression of Pldo^WT^. Note that such expression of Pldo^WT^ even creates a GOF phenotype, inducing the formation of filopodia-like protrusions. Rescued cell is marked by GFP (green) with MARCM technique. Scale bar corresponds to 10μm. C. Pldo^WT^ expression in hemocytes induce filopodia formation, similar to Cdc42^G12V^ expression, a known inducer of filopodia. Formation of filopodia in Cdc42^G12V^ background depends on *pldo* function (also see Suppl Figure S4D for quantification). Scale bar corresponds to 5μm. D-E. Filopodia formation induced by Pldo overexpression in hemocytes as driven by *hml-Gal4* driver (second panel from left, *GFP-IR* control, cf. to *wt* in left most panel) was suppressed by knock-down (via IR) of either *pldo, act, dia*, or *chic* (Profilin), and enhanced by *scar* knock-down. Scale bar corresponds to 5μm. (E) Quantification of number of filopodia per cell. At least 10 cells were quantified in each genotype in three independent experiments. Statistical analysis was 2-Way ANOVA, Tukey post-test with *** *P* < 0,0001.

Pldo overexpression in hemocytes induces a significant increase in filopodia-like protrusions formation (Figure 4C, left panel). To confirm that these protrusions were filopodia, we stained the cells with anti-Fascin, a known marker of mature filopodia (Vignjevic et al., 2006). We observed Fascin staining in some of the filopodia, in all such conditions (Suppl. Figure S4F). *cdc42* is a well-defined inducer of filopodia formation (Leung et al., 1998). As the Pldo induced protrusions resembled the effect of expressing constitutively active Cdc42 (Figure 4C, middle and left panel, respectively), we thus asked whether Cdc42^G12V^ induced filopodia in hemocytes depend on Pldo function. Strikingly, an RNAi knock-down of Pldo caused a total reversion of the Cdc42^G12V^ induced filopodia formation (Figure 4C, right panel, and Suppl. Figure S4C-D). Together, these data indicate that Pldo is essential for filopodia formation in hemocytes downstream of Cdc42. This result strongly support the notion that Pldo directly favors linear actin polymerization.

Similar to the rescue experiments of trichome formation in pupal wings, Pldo induced filopodia formation depends on regulators of linear actin polymerization, and as such its filopodia inducing effect was reversed by reducing the levels of Dia, Actin, or Chic/Profilin, and enhanced by reduction in the *scar* gene, promoting branched actin (Figure 4D-E). Again, this is consistent with the notion that Pldo promotes linear actin polymerization in the cellular competition for Actin monomers.

Taken together, the wing cuticle epithelial cell and hemocyte studies indicate that Pldo possesses a general function in regulating the competition for Actin monomers, favoring linear actin polymerization over branched polymerization.

### N-terminal region of Pldo is sufficient to induce filopodia formation

To evaluate which regions of the Pldo protein were essential for its function in actin polymerization, we generated a mutant isoform deleting its C-terminal portion (Pldo^ΔC^) (Figure 5A). We first compared the gain-of-function (GOF) phenotypes of Pldo^WT^ and Pldo^ΔC^ in the hemocyte assay, noting that both Pldo^*WT*^ and Pldo^ΔC^ induce filopodia formation equally well (Figure 5B-C and Suppl. Figure S5B).

**Figure 5:**
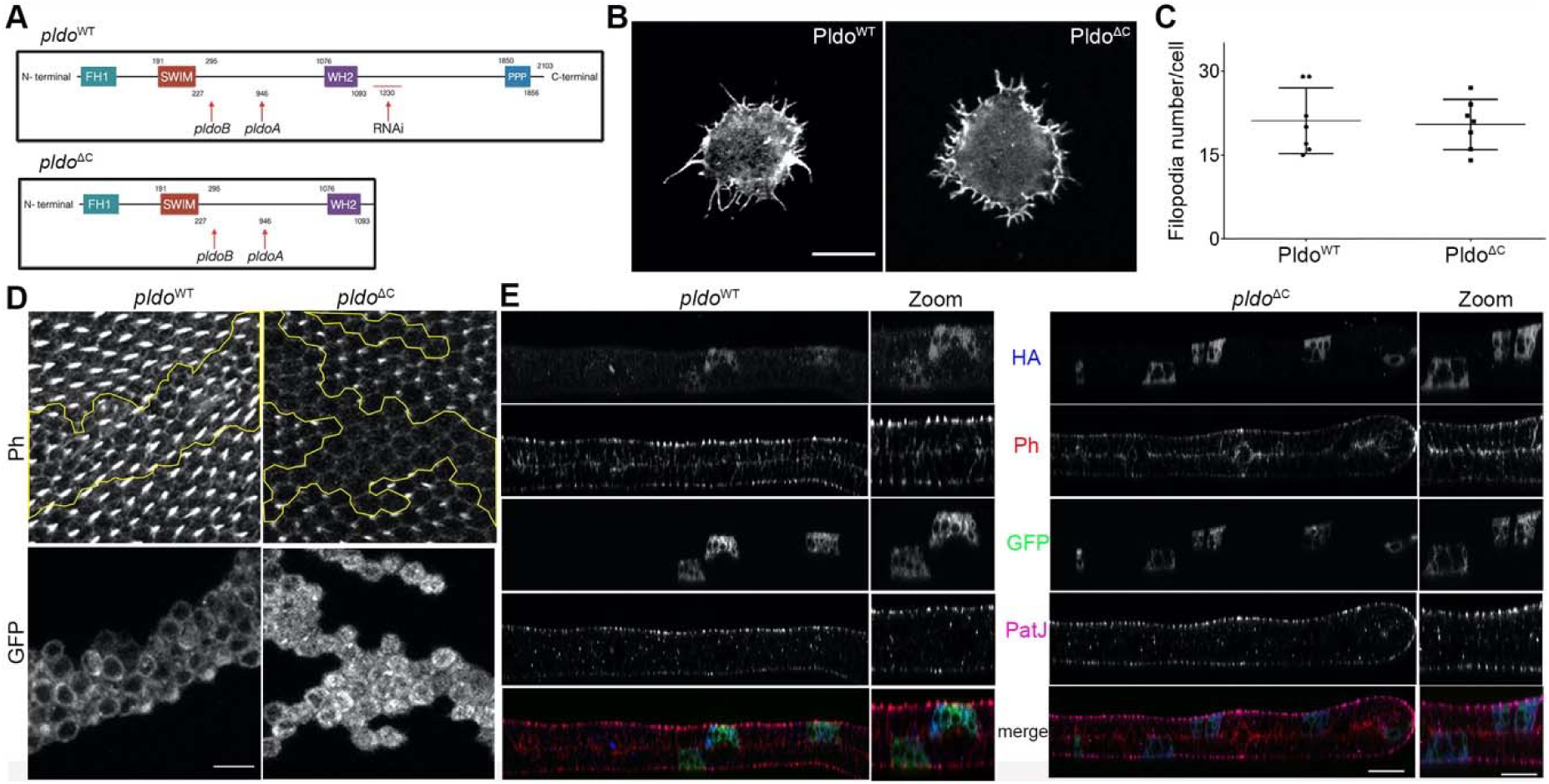
N-terminal region of Pldo is sufficient to induce filopodia formation. A. Schematic of Pldo highlighting deletion isoform with C-terminal deletion, Pldo^ΔC^ (note that the RNAi sequences are absent in the Pldo^ΔC^ construct). B-C. Pldo^ΔC^ was sufficient to induce formation of filopodia in hemocytes, to the same extent as full-length Pldo^WT^. Examples shown in (B) and quantification in (C), showing number of filopodia per cell. Scale bar represents 10 μm. At least 10 cells were analyzed in each condition in 3 independent experiments. Statistical analysis with t-test, *P*=0,8028, non-significant. D. Expression of Pldo^ΔC^ in MARCM *pldo* mutant cells (right panels) in pupal wings was not able to rescue hair formation, compare to *pldo*^WT^ expression (left panels). Distal is right. Scale bar represents 25 μm. E. Subcellular localization (z-plane) analyses of HA-tagged Pldo^WT^ (left panels) and Pldo^ΔC^ (right panels), displaying rhodamin-phalloidin (red) staining actin filaments, GFP (green) as clonal marker (via MARCM, mutant cells), PatJ (magenta) as a junctional (apical) cell marker, and HA showing Pldo localization (blue). Note that there is no detectable difference in the localization of Pldo^WT^ and Pldo^ΔC^ in pupal wing cells. Right panels (zoom) show higher magnification. Scale bar represents 25 μm.

In the context of actin hair formation in wing cells, however, Pldo^*WT*^ fully rescued the *pldo* LOF mutant phenotype, but expression of Pldo^ΔC^ in mutant clones did not rescue the loss of hair defect (Figure 5D). Similarly, Pldo^ΔC^ expression did not rescue the loss of hair phenotype in the *ptcGal4>pldo-IR* background (Suppl. Figure S5A). These results indicate that the N-terminal portion of Pldo is sufficient to induce linear actin polymerization, for example in hemocytes, but that its C-terminal region is required in more complex situations in epithelial cells, like trichome formation, likely due to a regulatory function or for subcellular complex formation. To test for the latter, we asked whether the C-terminal portion of Pldo is required to localize the protein to a specific region within the wing cells. Surprisingly, we did not detect a difference in the localization between Pldo^WT^ and Pldo^ΔC^ in either wing cells (Figure 5E and Suppl. Figure S5C-D), or hemocytes (Suppl. Figure S5B). Similar to Scar (Beli et al., 2008; Kunda et al., 2003; Rodriguez-Mesa et al., 2012; Zallen et al., 2002), Pldo localization is detected throughout the cell, which has also been noted for its human homologue ZSWIM8 (Okumura et al., 2021).

In this context, we also tried to identify the minimal portion of Pldo required to induce linear actin polymerization by generating additional C-terminal deletions (Pldo^1-685^ and Pldo^1-515^ isoforms, as compared to 1-1105 on Pldo^ΔC^). However, both constructs were cell lethal, in both *Drosophila* S2 cells and human A549 cells.

### Pldo function in actin cytoskeletal regulation is conserved in mammalian cells

We next wished to determine whether the function of Pldo in actin regulation is a conserved feature of the protein and used a cell migration assay in human cells to this end. Cell migration is important in many different developmental and physiological processes and requires specific actin cytoskeletal dynamics in a highly regulated process (Clainche and Carlier, 2008). Different cells use distinct mechanisms for migration and can also change between different types of migration. The common component in the distinct mechanisms of cell migration is Actin. Similarly, a key feature during cell migration is the upregulation of the Arp2/3 complex activity (Kelber and Klemke, 2011; Shellard and Mayor, 2020; Swaney, and Li, 2016). We thus used a cell migration assay as a tool for an initial insight whether Pldo function in actin regulation is conserved in human cells.

To this end, we used the human A549 cell line (kindly provided by the Bartel laboratory (Shi et al., 2020)). We established a clonal knock-out cell line for *ZSWIM8*^*-/-*^, with *ZSWIM8* being the human *pldo* orthologue, and analyzed its effect in a cell migration wound healing assay. Strikingly, cells mutant for *pldo/ZSWIM8* showed an increase in the migration rate, as compared to wild-type control A549 cells (Figure 6A-B). Importantly, this effect was fully reversed by the expression of *Drosophila* Pldo in those cells (Figure 6A-B), confirming that ZSWIM8/Pldo also regulates actin polymerization in mammalian cells and that the *Drosophila* Pldo protein is functionally equivalent in this context. These data are very similar to experiments demonstrating that branched actin filaments are essential in the wound healing assay (Zhao et al., 2020; using the same cell line), and strongly suggest that Pldo is also able to prevent branched actin polymerization.

**Figure 6:**
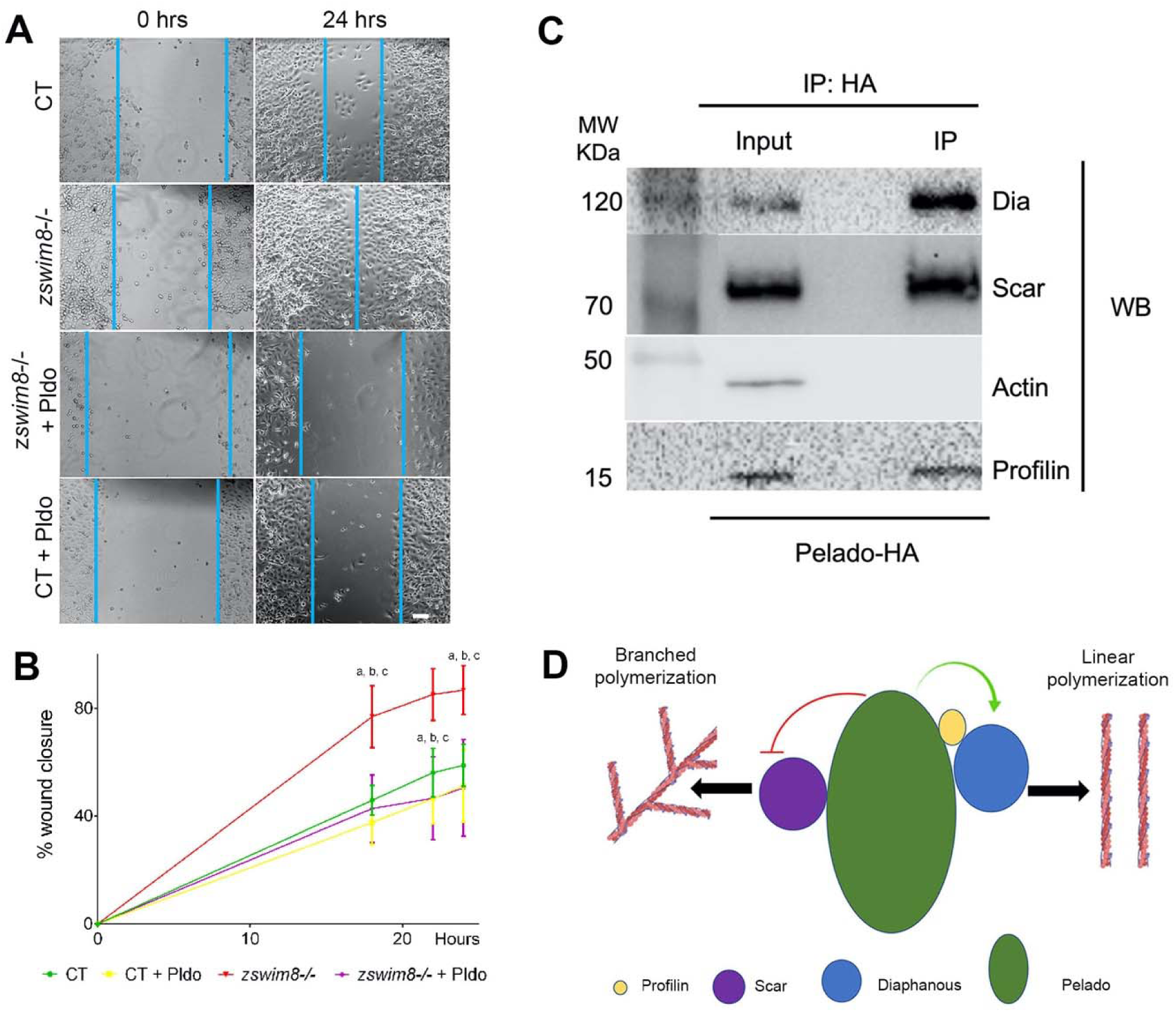
Pldo function in actin cytoskeletal regulation is conserved in human cells. A-B. Pldo/ZSWIM8 function was evaluated in a scratch wound migration assay in human cells (cell line A549). *pldo/ZSWIM8* mutant cells showed a significant increase in the rate of migration during assay. This effect was reversed by expressing the *Drosophila* Pldo gene, (independent experiments: n=3). Scale bar represents 50 μm. (B) Quantification of % wound closure over time. Statistical analysis was performed via a Two-Way ANOVA, Tukey post-test, *P*<0,0001, “a” indicates ** between CT and *zswim8-/-;* “b” indicates *** between CT+Pldo and *zswim8-/-;* and “c” indicates *** between *zswim8-/-* and *zswim8-/-* +Pldo. C. Immunoprecipitation (IP) assay indicating an interaction of Pldo with different proteins that regulate actin cytoskeletal dynamics in *Drosophila* S2 cells transfected with HA-Pldo^WT^ and immunoprecipitated with anti-HA. Note that Dia, Scar, and Chic/Profilin all did co-IP with Pldo, suggesting that they form a complex. Actin was not co-IPed with Pldo. A complete sample gel of this IP is shown in Suppl Figure S7. D. Schematic of proposed model with Pldo forming a multiprotein complex with Scar, Dia and Chic/Profilin during the regulation of actin cytoskeleton dynamics. Our data suggest that Pldo inhibits branched actin polymerization (probably through inhibiting Scar/WAVE activity) and promotes linear actin polymerization through Dia and Chic/Profilin.

These data strongly suggest that the Pldo/ZSWIM8 function in the regulation of actin cytoskeletal elements is conserved between *Drosophila* and human cells, and further confirm the notion that these proteins function in general to favor linear actin polymerization over branched filaments.

### Pldo physically associates with regulators of actin polymerization

The actin polymerization regulators mDia2 and Scar/WAVE are known to be part of the same molecular complex and that the binding of Cdc42 modulates their interaction to induce filopodia formation (Beli et al., 2008). We thus asked whether Pldo is also associated with these and/or other proteins that regulate actin cytoskeleton dynamics. To test this hypothesis, we used the *Drosophila* S2 R+ cell line (Schneider, 1972; which is a macrophage-like lineage derived from late embryos) and we observed that Pldo was indeed part of a molecular complex with Dia, Scar/WAVE, and Chic/Profilin (Figure 6C). These data suggest that Pldo could be regulating the activity of these proteins as part of the described complex (Beli et al., 2008), including Dia and Scar/WAVE.

Two reports characterizing EBAX-1, the *pldo* orthologues in *C. elegans*, and human ZSWIM8, respectively (Shi et al., 2020; Wang et al., 2013) suggest that it can function as an E3 Ubiquitin ligase, being part of a ubiquitin ligase complex, including Elongins B and C, and Cullin-2 (Wang et al., 2013). Since *pldo* LOF is reversed by reducing Scar levels, and Pldo and Scar are co-immunoprecipitated (Figure 6C, also Suppl. Figure S6A), we hypothesized that Scar could be a target of such a Pldo function, and thus if Scar were the target of Pldo in the context of a ubiquitin ligase complex, we would expect a change in Scar levels in mutant cells for *pldo*. We did not, however, observe an effect of *pldo* on Scar protein levels or localization, as compared to wild-type control cells (Suppl. Fig. 6B-C; see legend to Suppl Fig. S6 for more details).

In summary, our data is consistent with a model where Pldo is part of a multiprotein complex and it regulates the activity of its components in a manner that favors linear actin polymerization, either by promoting delivery of Actin monomers towards linear polymerization or by promoting Dia function or inhibiting Scar/WAVE (see model in Figure 6D).

## Discussion

Here we describe a lethal recessive mutation in the *pelado* (*pldo*) gene that, when analyzed in mosaic animals, results in external body regions lacking actin-based (hair) structures. Functional analyses of *Drosophila* Pelado (Pldo, a.k.a. ZSWIM8 in humans and other species) in the context of the regulation of actin polymerization dynamics indicate that it is an essential component required for linear actin polymerization. Our studies support a conserved function of Pldo/ZSWIM8 in promoting linear actin polymerization – at the expense of or in competition of branched polymerization - in several cell types, ranging from *Drosophila* epithelial cells to a human cell line. Pldo/ZSWIM8 proteins share multiple domains that have been implicated in the regulation of actin filament dynamics and Pldo/ZSWIM8 physically interacts with actin polymerization regulators, including Scar/WAVE, the formin Diaphanous, and Profilin. Functional dissection reveals specific domains of Pldo/ZSWIM8 being required for filopodia formation and other protein regions functioning in specific cellular contexts. Taken together, our data indicate that the N-terminal portion of Pldo/ZSWIM8 has a conserved regulatory function that is critical for linear actin polymerization. While ZSWIM8 has been linked to ubiquitination in some contexts (Shi et al., 2020; Wang et al., 2013), our data argue against such a requirement during the regulation of actin dynamics.

### Pldo/ZSWIM8 is required for linear actin polymerization

During development and morphogenesis *Drosophila* cuticular epithelial cells differentiate to produce the adult phenotypic adaptations as part of the exoskeleton. Wing development, leading to largely hexagonal cells with each cell producing an actin-based hair, serves as an excellent model to study the role of cellular morphogenesis and the regulation of cytoskeletal dynamics (Lu and Adler, 2015). The actin-based cellular hair, or trichome, covers all wing cells and most parts of the external body in *Drosophila* with the function of the trichome presumably related to a passive role directing airflow over the body and wing (Guild et al., 2005).

The positioning of the actin hairs, in the distal vertex of each cell, depends on the core Fz/PCP pathway and its effectors, which control the accumulation and polymerization of actin fibers at the apico-distal region of every cell (rev. in Adler, 2012; Carvajal-Gonzalez and Mlodzik, 2014; Strutt and Strutt, 2009). Seminal work revealed that hair formation is highly dependent on linear actin polymerization (Mitchell et al., 1983). Hair outgrowth in the wing epithelia has been divided in three different stages: at 28 hours APF Actin starts to accumulate in the apico-distal zone of wing cells, aided by tubulin dependent transport and vesicle recycling (Gault et al., 2012), directed by the activity of the core PCP pathway; at 30-32 hours APF, the prehair structure becomes apparent corresponding to the early stages of hair growth, which is focused in one distal spot due to the inhibitory activity of Mwh (Adler, 2012; Carvajal-Gonzalez and Mlodzik, 2014). Subsequently, at 32-36 hours APF, the hairs extend and are completely formed (at 36h APF), mainly by elongation of linear actin filaments, arising from the “prehair” foci. A detailed analysis of *pldo/zswim8* mutants during the different stages of hair growth suggests that it is required for the formation of the prehair and linear actin extension within the hair. In addition, *pldo/zswim8* appears to affect cell shape morphogenesis with a reduction in hexagonal cell shapes, consistent with a role in linear cortical actin filaments. The observation that overexpression of the formin Dia can rescue the loss-of-function (LOF) *pldo/zswim8* phenotype resulting in loss of hairs, supports the notion that Pldo/ZSWIM8 regulates formin-dependent linear actin polymerization. The other described genes whose LOF generates the absence of cellular hairs in *Drosophila* are *shavenbaby* (*svb*) and *shavenoid* (*sha*). Svb is a transcription factor that induces the expression of Sha, which has been defined as a protein that regulates the actin cytoskeleton. However, *sha* is not conserved in the animal kingdom and its presence is specific to insects and arthropods (He and Adler, 2002; www.flybase.org).

Actin hair formation, both number and location, is regulated by the core Fz/PCP pathway, but little is known about the mechanisms that regulate the actual actin polymerization during this process. Core PCP factor polarization, as exemplified by Fmi, is not affected in *pldo/zswim8* mutant cells, suggesting that Pldo/ZSWIM8 is a downstream effector of the PCP pathway. Dsh accumulates at the apico-distal zone of *Drosophila* wing cells, where it is thought to recruit and activate RhoA, Rac, and Daam1 (Disheveled-associated activator of morphogenesis 1, originally identified in Xenopus (Habas et al., 2001). Daam1 and likely also other formins, like Dia (as Daam1 appears to be redundant in this wing hair context), interact with Profilin to induce actin polymerization (Adler, 2012; Barko et al., 2010; Goodrich and Strutt, 2011; Matusek et al., 2008; Singh and Mlodzik, 2012; Yaniv et al., 2020). The *pldo/zswim8* LOF phenotype is rescued by Dia or Profilin overexpression, while different isoforms of dDaam1, including activated ones (Matusek et al., 2006), do not rescue *pldo/zswim8* LOF defects, consistent with the notion that Dia and not Daam1 is the critical formin family member in this context. As an association of Dia and the Dsh complex has not yet been reported, the connection between Pldo/ZSWIM8 and core PCP complexes remains unclear and further work will be required to establish a better understanding of the information flow of the regulatory interactions.

Besides the actin hair/trichome, *pldo/zswim8* mutant epithelial cells also display cell shape defects, which – based on its link to actin – are likely caused by defects in cortical actin regulation. As the cortical actin cytoskeleton fulfills an essential role in all cells, it is not surprising that a *pldo/zswim8* mutant is embryonic lethal, albeit with a very variable phenotype. It should be noted that the highest levels of *pldo/zswim8* message are maternal in the early embryo, but germline clones of *pldo/zswim8* do not survive to allow egg development and deposition, likely due to defects during oogenesis.

### Pldo/ZSWIM8 affects the balance of linear versus branched actin filament polymerization

The loss of the actin hair observed in *pldo/zswim8* mutant cells is not only rescued by the overexpression of the formin Dia and Actin itself, but also by the knockdown of *scar* and *arp2/3*. These ‘synthetic’ genetic interactions suggest that Pldo/ZSWIM8 affects the balance of linear and branched actin polymerization, with either an excess of Actin monomers or the reduction of the capability of the cell to promote branched Actin changes the balance back to linear Actin. Thus, Pldo/ZSWIM8 facilitates the process by favoring linear polymerization. Notably, *wasp* knockdown did not reverse the *pldo/zswim8* mutant phenotype, as compared to *scar*, which efficiently reversed the *pldo/zswim8* defect. It has been described that Scar and Wasp act independently, regulating branched actin polymerization in different development contexts, as for example, Scar is important for morphogenetic cell processes, while Wasp being essential during muscle and sensory organ development (Brüser and Bogdan, 2015). Interestingly, a knock-down of *mwh*, an anti-formin inhibiting Dia, allowed also a partial phenotypic reversion of the *pldo/zswim8* defect, suggesting that Dia function is rate limiting in *pldo/zswim8* mutants (Mwh is actually epistatic to Pldo/ZSWIM8 as the *mwh* LOF hair phenotype is manifested in the double know-down, with 3+ hairs forming per cell (Yan et al., 2008)).

Taken together, Pldo/ZSWIM8 serves a balance function promoting linear actin polymerization, which is mediated by Dia, as a formin, inducing linear actin polymerization, in competition with Scar, inducing branched actin polymerization. Therefore, the balance between linear and branched actin polymerization appears to be a key determinant in the trichome formation process. This effect of Pldo/ZSWIM8 may be comparable to the one described for Profilin (Suarez et al., 2015), which favors linear actin polymerization over branched filaments in fission yeast cells. Here, in the absence of Profilin new Arp2/3 complex mediated branched filaments are continuously generated, while Dia2 assembled linear actin filaments elongate 4-times slower in the absence of Profilin, as compared to control filaments (Suarez et al., 2015).

The notion that Pldo/ZSWIM8 favors linear actin polymerization is also supported by the phenotype observed in hemocytes. Here, *pldo/zswim8* mutant and Pldo/ZSWIM8 overexpression backgrounds generate cell morphology changes involving rearrangements in actin-dependent structures, like filopodia and lamellipodia. *pldo/zswim8* mutants cells display a rounded morphology with less surface contacts and increased ruffles, structures mainly based on branched actin polymerization (Campellone and Welch, 2010; Legg et al., 2007). Similar observations were made for Scar gain-of-function (GOF) scenarios in BG3 cells from *Drosophila* (Cloud et al., 2019). It is noteworthy that cell adhesion to the matrix is based in focal adhesion related structures, with focal adhesions directly interacting with the substrate and being associated with linear actin filaments, known as stress fibers (Hohmann and Dehghani, 2019a; Kelber and Klemke, 2011). These observations in hemocytes reinforce the idea that Pldo/ZSWIM8 functions to promote linear actin polymerization. Accordingly, Pldo/ZSWIM8 overexpression in hemocytes induces the formation of filopodia, structures essentially formed by long linear actin filaments. Importantly, Pldo/ZSWIM8 LOF in hemocytes expressing the constitute active form of Cdc42, prevented filopodia formation, strongly suggesting that Pldo/SWIM8 has a role in lineal actin polymerization. As such, the actin hair outgrowth process in epithelial cells is highly similar to filopodia formation, with both being protrusions of the plasma membrane driven by linear actin filament extension/growth. Similar to the epithelial actin hair context, *pldo/zswim8* mutant defects in hemocytes are reversed by enhancing linear actin polymerization either by (i) increasing formin activity by overexpressing Dia, (ii) reducing branched polymerization by knocking-down Scar, or (iii) by increasing Actin monomer availability via overexpressing Actin itself.

Importantly, this function of Pldo/ZSWIM8 is conserved from *Drosophila* to mammals. In a well-established wound healing cell migration assay in the human A549 cell line (Han et al., 2016; Kim et al., 2020; Liao and Peng, 2020; de Oliveira Rodrigues et al., 2020; Piao et al., 2014; Wang et al., 2017; Zhao et al., 2020), we observed that *pldo/zswim8* knockout cells migrate significantly faster than control cells, suggesting that these cells have an increase in branched actin polymerization, allowing a faster migration. Strikingly, this effect is rescued by providing *Drosophila* Pldo/ZSWIM8 protein to these knockout human A549 cells. This not only suggests a conserved function, but also confirms that the *Drosophila* factor is functionally equivalent to the human protein. Very similar results were described by knocking down WDR63, an inhibitor of the Arp2/3 complex, causing an increase in branched polymerization and cell migration in A549 cells (Zhao et al., 2020), opposite to the effect observed when *arp2* is knocked down. These results are fully consistent with our data in *Drosophila* contexts, indicating that Pldo/ZSWIM8 changes the balance between linear and branched actin polymerization. This might be mediated possibly by either reducing Scar/WAVE activity (and in consequence Arp2/3 dependent branched actin nucleation) or by promoting Dia/formin function and hence linear actin polymerization, or both.

In summary, using multiple different cell types and experimental strategies, we have demonstrated that Pldo/ZSWIM8 serves a conserved function to promote linear actin polymerization at the expense of branched actin. Our results are all consistent with the notion that Pldo/ZSWIM8 functions in the same direction as Dia and Profilin/Chic activity and opposed to Scar and Arp2/3 activities in promoting linear actin polymerization. In this context, Pldo/ZSWIM8 seems to favor Actin monomer competition and recruitment towards linear filament polymerization, since Actin overexpression or knock-down can reverse the respective phenotypes induced in wing epithelial cells and hemocytes by Pldo/ZSWIM8 knock-down or overexpression, respectively. Altogether, our results suggest that Pldo/ZSWIM8 is able to induce linear actin polymerization (as is required for filopodia formation in hemocytes, for example) and to prevent branched actin polymerization (as shown in *pldo/zswim8* LOF experiments in hemocytes and human cells).

### Pldo/ZSWIM8 as a part of a molecular complex

mDia2, Arp2/3, and Scar/WAVE are detected in a multimolecular complex in HeLa cells, and these interactions are modulated by Cdc42 (Beli et al., 2008). We tested for co-immunoprecipitation of Pldo/ZSWIM8 with different proteins that regulate actin filament dynamics. Pldo/ZSWIM8 appears to be indeed part of a molecular complex with Dia, Scar, and Profilin, suggesting that Pldo/ZSWIM8 is part of a similar molecular complex to the one described (Beli et al., 2008).

ZSWIM8 (EBAX-1 in *C. elegans*) has also been characterized as an E3 ligase and a component of a ubiquitin ligase complex (Shi et al., 2020; Wang et al., 2013), and thus we asked whether Pldo/ZSWIM8 could have the same function in the context of the regulation of actin filament dynamics. Since a *scar* knockdown reversed the *pldo/zswim8* LOF phenotype, we tested whether Scar could be a potential degradation target of Pldo/ZSWIM8 in the actin context. We did not detect any changes to Scar levels, however, despite the fact that Scar and Pldo/ZSWIM8 are found in the same molecular complex. Nonetheless, although we did not detect any evidence for a ubiquitin-mediated degradation function of Pldo/ZSWIM8 in the actin cytoskeleton regulation context, we cannot rule out that Pldo/ZSWIM8 might have a function as part of a ubiquitin ligase complex in this context as well. Pldo/ZSWIM8 might have distinct molecular functions in different contexts. It will be interesting to determine potential regulatory conditions and mechanisms that select one function of Pldo/ZSWIM8 over the other, or if there is a cell-type or stage specificity to potential distinct molecular functions of this interesting protein family.

### Pldo/ZSWIM8 and a potential role in human disease

As suggested by Yamamoto, Bellen and colleagues (Yamamoto et al., 2014), Pldo/ZSWIM8 appears to be linked to a genetic disorder in humans affecting the nervous system. This observation could be explained by our data regarding its function in the regulation of actin cytoskeletal dynamics, as biogenesis of neurites requires a highly regulated actin polymerization network (Ben-Yaacov et al., 2001; Korobova and Svitkina, 2008; Matusek et al., 2008; Santos et al., 2002; Tahirovic et al., 2010; Yaniv et al., 2020). For example, linear actin polymerization is required for radial glial migration and in the initial stages of neurite formation and branched polymerization is essential for their maturation. As such, Pldo/ZSWIM8 levels, when altered, could easily affect this process leading to neurological defects. It is also important to consider that there are five isoforms of Pldo/ZSWIM8 in humans, making it more difficult to analyze. In addition, taken together with the data presented here, Pldo/ZSWIM8 could also have a role in cancer cells, since its function might modify the migratory capacity of invasive metastatic cells.

## Materials and Methods

### *Drosophila melanogaster* strains and phenotypic analysis in adult wings

*Drosophila* flies were grown on standard food at 25ºC, and phenotypes were analyzed at this temperature, unless otherwise indicated.

The following fly strains were used:

Bloomington (BL): BL5137 UAS-CD8::GFP,

BL56751 UAS-Diaphanous,

BL7310 UAS-Actin,

BL18553 UAS-dsRNA *pldo*,

BL52333 *pldo*^*A*^,

BL52334 *pldo*^*B*^,

BL41552 UAS-dsRNA GFP,

BL1709 FRT19A,

BL34523 UAS-dsRNA *chic*,

BL5905 *w*^*1118*^,

BL24651 UAS-Dicer^2^,

BL4854 UAS-Cdc42^G12V^.

VDRC strains (all are UAS-dsRNA constructs):

21908 *scar*, 13759 *wasp*, 41514 *mwh*, 102759 *chic*, 27623 *quail*, 101438 *actin UAS Scar-GFP* was a gift from S. Bogdan (Stephan et al., 2011).

Overexpression of cDNA transgenes or RNAi (IR) was performed using the Gal4/UAS system (Brand and Perrimon, 1993). The Gal4 expression drivers used were as follows: *patched-Gal4, Hml-Gal4, pannier-Gal4*, and *ey-Gal4*. Where indicated, UAS-Dicer2 was included with *UAS-IR* expression to increase RNAi efficiency.

All stocks employed in experiments were generated by standard crosses.

#### MARCM analysis

For pupal wing clone generation, offspring were subjected to heat shock (at 37ºC) for 1 hour at 80h +/-12h after egg laying. Pupal clones were thus marked by GFP expression. Third instar larvae possessing clones (GFP positive) were selected and determined the 0 time (white prepupae) to proceed with immunofluorescence stainings at specific times during animal development (and hair formation).

In hemocytes, the offspring were subjected to heat shock at 48h, 72h and 96h after egg laying for 1 hour, then incubated at 18ºC for 1 hour.

For analysis of adult wing trichomes, adult wings were removed, incubated in wash buffer (PBS and 0.1% Triton X-100), and mounted on a slide in 80% glycerol in PBS.

In the trichome rescue assay, classification of the phenotypic severity of absence of hairs in adult wings was performed by visual inspection. Criteria for classification were the extent of area without hairs. Phenotypes were classified as rescued (no area without hairs), partial rescue (small area without hairs) and no rescue (large area without hairs).

To analyze trichomes in adult nota (dorsal thorax), flies were partially dissected, incubated at 95ºC in 10% KOH for 10 min to clear fat tissue, washed (PBS and 0.1% Triton X-100) and the placed in 80% glycerol in PBS. Nota were fully dissected and mounted on a slide in 80% glycerol in PBS. Adult wings and nota were imaged at room temperature on a Axioplan (Carl Zeiss) microscope. Images were acquired with a Zeiss AxioCam (Color type 412-312) and AxioCam software.

### Sequence analysis

Sequence analysis and predictions were made at online programs Psipred and ELM, the eukaryotic linear motif resource for functional sites in proteins.

### Immunofluorescence

For analysis in pupal wings, white pupae were collected (0 h APF) and aged at 25ºC until dissection. Dissections were performed as follows: in brief, pupae were immobilized on double-sided tape, removed from the pupal case, and placed into PBS, in which pupae were partially dissected to remove fat tissue, fixed in 4% paraformaldehyde in PBS for 45 minutes at room temperature (RT) or overnight at 4ºC, and washed 3 x in PBS and 0.1% Triton X-100. Wing membranes were removed, and immunostaining was performed by standard techniques. In brief, tissue was incubated in wash buffer containing 10% normal goat serum overnight for primary antibody (4ºC), washed 3 x with PBS, and incubated with secondary antibody for at least 2 h (25ºC) and fluorescent phalloidin for staining actin cytoskeleton. Wings were washed 3x with PBS and mounted in Mowiol. Pupal wing images were acquired at RT using a confocal microscope, either Zeiss LSM 710 or Leica SP8.

Images were processed with ImageJ (National Institute of Health).

For hemocyte immunostaining, primary culture was performed as described (Tirouvanziam et al., 2004). Briefly, third instar larvae were washed and a small incision on the posterior side of larvae was made to obtain hemocytes, which were incubated for 1h 15 min, fixed and stained. Samples were maintained in Vectashield until pictures were taken. Confocal images were captured using either a Zeiss LSM 710 or a Leica SP8 confocal microscopes. Images were processed using ImageJ.

### Antibodies

The following antibodies were used:

mouse anti-Fmi (1:10, Developmental Studies Hybridoma Bank),

mouse anti-Fascin (1:10, Developmental Studies Hybridoma Bank),

mouse anti-Scar (1:50, Developmental Studies Hybridoma Bank),

rabbit anti-Dia (1:1000, donated by P. Adler),

mouse anti-Chic (1:10, Developmental Studies Hybridoma Bank),

rabbit anti-PatJ (1:500), mouse anti-HA (1:50, Biologend),

mouse anti-β-actin (1:1000, Sigma).

All fluorophor-conjugated secondary antibodies were used at 1:200 and obtained from Jackson ImmunoResearch Laboratories, Inc.

Rhodamine-phalloidin was from Invitrogen and used at 1:400.

### Image analysis

Individual channels from color images were converted to grey scale using ImageJ. Fluorescent intensity levels were measured on maximal projections of image stacks using ImageJ.

### DNA construct generation

Pelado ΔC was generated by PCR using the following primers: Fw: 5’ CAC CAT GGA CCG CTT CAG CTT CG 3’, Rv 1: 5’ CTC TCC TCG CGT CTT AAA GGT TCC 3’, RV 2: 5’ GAA AAT AAA CGT GGC CAG GTT AAT AGG CAA TAG 3’, RV 3: 5’ GCT GAG TGT AAC TAG GGC ATC GAA AAT AAA CGT GGC 3’. PCR product was purified and cloned into pENTR vector, sequenced and using the Gateway kit was cloned into the pUASt vector, with the coding sequence of HA tag at the 5’ of *pldo* gene. Transgenesis was performed at BestGene. The deletions Pldo^1-685^ and Pldo^1-515^ were generated by restriction enzymes KflI and BsiWI, respectively. To express Pldo in mammalian cells, the coding sequence was cloned into VVPW/BC vector (donated by L. Gusella), using XbaI and EcoRI restriction sites.

### Cell culture, transfection, and scratch wound healing assay

*Drosophila* S2R+ cells were obtained directly from Drosophila Genomics Resource Center (DGRC), Indiana. Cells were grown in Schneider’s medium, supplemented with 10% FBS and maintained at 25ºC, and transfected with Effectene reagent, following standard protocols. Cells were lysed or immunostained 48 h after transfection in lysis buffer.

Human A549 cells were obtained from the Bartel Laboratory and were grown in DMEM medium, supplemented with 10% FBS and Puromycin (10 μg/μL) and maintained in 5% CO_2_ at 37ºC. Monoclonal cell line was generated by seeding individual cells in a 96-well plate, monitoring cells for growth, identifying individual colonies, and expanding monoclonal lines of interest. Clonal cells knock-out for *pldo/zswim8* were identified by PCR with the primers described (Shi et al., 2020). Cells were transfected with Effectene reagent, following standard protocols.

Cell cultures were plated to a confluence of 50-70%, and transfected the next day. After 8-12 h the medium was changed for fresh medium. When confluence was reached the scratch was performed with a yellow plastic tip. Pictures were taken at times 0, 18, 22 and 24 hours after the scratch was made.

### Western blotting and immunoprecipitation

S2R+ cells (grown in Schneider medium using standard procedures) were transfected with HA-Pldo and GFP-Scar, using Effectene reagent. After 48h protein extraction was performed. Immuno-precipitation (IP) and co-IP experiments were performed by incubating lysates with anti-HA or anti-GFP antibody, at 4ºC overnight followed by Agarose-A beads incubation, 5x washing steps, elution in SDS sample buffer. Samples were boiled at 95ºC for 10 minutes and proceeded to Western blotting.

### Statistical Analyses

Quantifications were made in ImageJ. To determine statistical differences between data sets of continuous data, we performed non-parametric ANOVA or t-test analyses on GraphPad. Specific analysis is indicated each time and *P* values lower than 0,05 were considered significant. Significance is indicated by asterisks. Sample sizes are described for each experiment in the figure legends. All experiments were performed on a minimum of 3 animals per condition. The statistical parameters are reported in figure legends. *N* indicates the number of animals and the error bars represent the standard error of the mean (SEM).

## Acknowledgements

We thank Sven Bogdan, David Bartel, Luca Gusella, and Steven Wassarman for generously sharing *Drosophila* strains, cell lines, and other reagents. We are most grateful to Luca Gusella for support and sharing equipment during the cell culture experiments. We gratefully acknowledge the Bloomington Drosophila Stock Center, and the Vienna Drosophila RNAi Center for supplying fly stocks. We thank all Mlodzik lab members for helpful discussions and advice. This work was supported by grant R35 GM127103 from the NIGMS/NIH to MM; FONDECYT grant 1190119, ANID and the Center of Genome Regulation, Fondap 15200002, ANID to AG, and Biomedical Neuroscience Institute, Iniciativa Científica Milenio ICM P09-015F to PO. CMP was in part supported by a National PhD scholarship number 21140101, CONYCIT/ANID.

## Author contributions

CMP, PO, MM, and AG conceptualized the project and designed the study and wrote the paper. AG identified the gene, and CMP and PO performed the experiments. PO, AG, and MM secured funding for the project and guided its completion.

## References

Adler, P.N. (2012). The frizzled/stan pathway and planar cell polarity in the Drosophila wing. Curr. Top. Dev. Biol. 101, 1–31.

Adler, P.N. (2018). A cytoskeletal activator and inhibitor are downstream targets of the frizzled/starry night planar cell polarity pathway in the Drosophila epidermis. Prog. Biophys. Mol. Biol. 137, 69–75.

Adler, P.N., Sobala, L.F., Thom, D., and Nagaraj, R. (2013). Dusky-like is required to maintain the integrity and planar cell polarity of hairs during the development of the Drosophila wing. Dev. Biol. 379, 76–91.

Aigouy, B., Farhadifar, R., Staple, D.B., Sagner, A., Röper, J.C., Jülicher, F., and Eaton, S. (2010). Cell Flow Reorients the Axis of Planar Polarity in the Wing Epithelium of Drosophila. Cell 142, 773–786.

Barko, S., Bugyi, B., Carlier, M.F., Gombos, R., Matusek, T., Mihályand, J., and Nyitrai, M. (2010). Characterization of the biochemical properties and biological function of the formin homology domains of Drosophila DAAM. J. Biol. Chem. 285, 13154–13169.

Beli, P., Mascheroni, D., Xu, D., and Innocenti, M. (2008). WAVE and Arp2/3 jointly inhibit filopodium formation by entering into a complex with mDia2. Nat. Cell Biol. 10, 849–857.

Ben-Yaacov, S., Le Borgne, R., Abramson, I., Schweisguth, F., and Schejter, E.D. (2001). Wasp, the Drosophila Wiskott-Aldrich syndrome gene homologue, is required for cell fate decisions mediated by Notch signaling. J. Cell Biol. 152, 1–13.

Berger, S., Schäfer, G., Kesper, D.A., Holz, A., Eriksson, T., Palmer, R.H., Beck, L., Klämbt, C., Renkawitz-pohl, R., and Önel, S. (2008). WASP and SCAR have distinct roles in activating the Arp2/3 complex during myoblast fusion. J. Cell Sci. 121, 1303–1313.

Bogdan, S., Schultz, J., Grosshans, J., Neurobiologie, I., Münster, U., Würzburg, U., and Biochemie, I. (2013). Physiological roles of Diaphanous (Dia) in actin dynamics Formin ‘ cellular structures. Commun. Integr. Biol. 1–12.

Brand, A.H., and Perrimon, N. (1993). Targeted gene expression as a means of altering cell fates and generating dominant phenotypes. Development 118, 401–415.

Brüser, L., and Bogdan, S. (2015). Molecular Control of Actin Dynamics In Vivo□: Insights from Drosophila. Handb. Exp. Pharmacol. 235, 251–263.

Burke, T.A., Christensen, J.R., Barone, E., Suarez, C., Sirotkin, V., and Kovar, D.R. (2014). Report Homeostatic Actin Cytoskeleton Networks Are Regulated by Assembly Factor Competition for Monomers. Curr. Biol. 24, 579–585.

Campellone, K.G., and Welch, M.D. (2010). A nucleator arms race: cellular control of actin assembly. Nat. Rev. Mol. Cell Biol. 11, 237–251.

Carlier, M.F., and Shekhar, S. (2017). Global treadmilling coordinates actin turnover and controls the size of actin networks. Nat. Rev. Mol. Cell Biol. 18, 389–401.

Carlier, M., Pernier, J., and Avvaru, B.S. (2013). Control of Actin Filament Dynamics at Barbed Ends by WH2 Domains: From Capping to Permissive and Processive Assembly. Cytoskeleton 70, 540–549.

Carvajal-Gonzalez, J.M., and Mlodzik, M. (2014). Mechanisms of planar cell polarity establishment in Drosophila. F1000Prime Rep. 6, 1–10.

Carvajal-Gonzalez, J.M., Mulero-Navarro, S., Smith, M., and Mlodzik, M. (2016). A novel frizzled-based screening tool identifies genetic modifiers of planar cell polarity in drosophila wings. G3 Genes, Genomes, Genet. 6, 3963–3973.

Chen, X.J., Squarr, A.J., Stephan, R., Chen, B., Higgins, T.E., Barry, D.J., Martin, M.C., Rosen, M.K., Bogdan, S., and Way, M. (2014). Article Ena / VASP Proteins Cooperate with the WAVE Complex to Regulate the Actin Cytoskeleton. Dev. Cell 30, 569–584.

Chou, F.-S., and Wang, P.-S. (2016). The Arp2/3 complex is essential at multiple stages of neural development. Neurogenesis 3, e1261653.

Clainche, C.L.E., and Carlier, M. (2008). Regulation of Actin Assembly Associated With Protrusion and Adhesion in Cell Migration. Physiol. Rev. 88, 489–513.

Classen, A., Aigouy, B., Giangrande, A., and Eaton, S. (2008). Imaging Drosophila Pupal Wing Morphogenesis. In Methods in Molecular Biology: Drosophila: Methods and Protocols, pp. 265–275.

Cloud, V., Thapa, A., Morales-Sosa, P., Miller, T.M., Miller, S.A., Holsapple, D., Gerhart, P.M., Momtahan, E., Jack, J.L., Leiva, E., et al. (2019). Ataxin-7 and non-stop coordinate scar protein levels, subcellular localization, and actin cytoskeleton organization. Elife 8, 1–35.

Cooper, J.A. (2013). Mechanisms of cell migration in the nervous system. J. Cell Biol. 202, 725–734.

Cruz, C., Glavic, A., Casado, M., and De Celis, J.F. (2009). A gain-of-function screen identifying genes required for growth and pattern formation of the Drosophila melanogaster wing. Genetics 183, 1005–1026.

Davidson, A.J., and Wood, W. (2016). Unravelling the Actin Cytoskeleton: A New Competitive Edge? Trends Cell Biol. 26, 569–576.

Davidson, A.J., Amato, C., Thomason, P.A., and Insall, R.H. (2018). WASP family proteins and formins compete in pseudopod- and bleb-based migration. J. Cell Biol. 217, 701–714.

Delon, I., Chanut-Delalande, H., and Payre, F. (2003). The Ovo / Shavenbaby transcription factor specifies actin remodelling during epidermal differentiation in Drosophila. Mech. Dev. 120, 747–758.

Eaton, S., Wepf, R., and Simons, K. (1996). Roles for Racl and Cdc42 in Planar Polarization and Hair Outgrowth in the Wing of Drosophila. J. Cell Biol. 135, 1277–1289.

Etournay, R., Popović, M., Merkel, M., Nandi, A., Blasse, C., Aigouy, B., Brandl, H., Myers, G., Salbreux, G., Jülicher, F., et al. (2015). Interplay of cell dynamics and epithelial tension during morphogenesis of the Drosophila pupal wing. Elife 4, 1–51.

Fagan, J.K., Dollar, G., Lu, Q., Barnett, A., Jorge, J.P., Schlosser, A., Pfleger, C., Adler, P., and Jenny, A. (2014). Combover/CG10732, a Novel PCP Effector for Drosophila Wing Hair Formation. PLoS One 9, e107311.

Faix, J., and Grosse, R. (2006). Staying in Shape with Formins Review. Dev. Cell 10, 693–706.

Fang, X., and Adler, P.N. (2010). Regulation of cell shape, wing hair initiation and the actin cytoskeleton by Trc/Fry and Wts/Mats complexes. Dev. Biol. 341, 360–374.

Fang, X., Lu, Q., Emoto, K., and Adler, P.N. (2010). The drosophila fry protein interacts with Trc and is highly mobile in vivo. BMC Dev. Biol. 10.

Fletcher, D. a, and Mullins, R.D. (2010). Cell mechanisms and cytoskeleton. Nature 463, 485–492.

Frankel, S., and Moosekert, M.S. (1994). The actin-related proteins. Yeast 70, 30–37.

Fricke, R., Gohl, C., Dharmalingam, E., Grevelhorster, A., Zahedi, B., Harden, N., Kessels, M., Qualmann, B., and Bogdan, S. (2009). Drosophila Cip4 / Toca-1 Integrates Membrane Trafficking and Actin Dynamics through WASP and SCAR / WAVE. Curr. Biol. 19, 1429–1437.

Gao, B. (2012). Wnt Regulation of Planar Cell Polarity (PCP). Curr. Top. Dev. Biol. 101, 263–295.

Gault, W.J., Olguin, P., Weber, U., and Mlodzik, M. (2012). Drosophila CK1-G, gilgamesh, controls PCP-mediated morphogenesis through regulation of vesicle trafficking. JCB 196, 605–621.

Gautreau, A.M., Fregoso, F.E., Simanov, G., and Dominguez, R. (2021). Nucleation, stabilization, and disassembly of branched actin networks. Trends Cell Biol. 1–12.

Godin, S.K., Meslin, C., Kabbinavar, F., Bratton-Palmer, D.S., Hornack, C., Mihalevic, M.J., Yoshida, K., Sullivan, M., Clark, N.L., and Bernstein, K.A. (2015). Evolutionary and functional analysis of the invariant SWIM domain in the conserved Shu2/SWS1 protein family from Saccharomyces cerevisiae to Homo sapiens. Genetics 199, 1023–1033.

Goley, E.D., and Welch, M.D. (2006). The ARP2/3 complex: an actin nucleator comes of age. Nat. Rev. Mol. Cell Biol. 7, 713–726.

Goodrich, L. V., and Strutt, D. (2011). Principles of planar polarity in animal development. Development 138, 1877–1892.

Guild, G.M., Connelly, P.S., Ruggiero, L., Vranich, K.A., and Tilney, L.G. (2005). Actin Filament Bundles in Drosophila Wing Hairs□: Hairs and Bristles Use Different Strategies for Assembly. Mol. Biol. Cell 16, 3620–3631.

Gurung, R., Yadav, R., Brungardt, J.G., Orlova, A., Egelman, E.H., and Beck, M.R. (2016). Actin polymerization is stimulated by actin cross-linking protein palladin. Biochem. J. 473, 383–396.

Habas, R., Kato, Y., and He, X. (2001). Wnt/Frizzled activation of Rho regulates vertebrate gastrulation and requires a novel formin homology protein Daam1. Cell 107, 843–854.

Han, M.L., Zhao, Y.F., Tan, C.H., Xiong, Y.J., Wang, W.J., Wu, F., Fei, Y., Wang, L., and Liang, Z.Q. (2016). Cathepsin L upregulation-induced EMT phenotype is associated with the acquisition of cisplatin or paclitaxel resistance in A549 cells. Acta Pharmacol. Sin. 37, 1606–1622.

He, B., and Adler, P.N. (2002). The genetic control of arista lateral morphogenesis in Drosophila. Dev Genes Evol 212, 218–229.

He, Y., Fang, X., Emoto, K., Jan, Y., and Adler, P.N. (2005). The Tricornered Ser / Thr Protein Kinase Is Regulated by Phosphorylation and Interacts with Furry during Drosophila Wing Hair Development. Mol. Biol. Cell 16, 689–700.

Henson, J.H., Yeterian, M., Weeks, R.M., Medrano, A.E., Brown, B.L., Geist, H., Pais, M., Oldenbourg, R., and Shuster, C. (2015). Arp2 / 3 complex inhibition radically alters lamellipodial actin architecture, suspended cell shape, and the cell spreading process. Mol. Biol. Cell 26, 887–900.

Hohmann, and Dehghani (2019a). The Cytoskeleton—A Complex Interacting Meshwork. Cells 8, 362.

Hohmann, T., and Dehghani, F. (2019b). The cytoskeleton-a complex interacting meshwork. Cells 8, 1–58.

Ito, T., Narita, A., Hirayama, T., Taki, M., Iyoshi, S., Yamamoto, Y., Maéda, Y., and Oda, T. (2011). Human Spire Interacts with the Barbed End of the Actin Filament. J. Mol. Biol. 408, 18–25.

Jaiswal, R., Stepanik, V., Rankova, A., Molinar, O., Goode, B.L., and Mccartney, B.M. (2013). Drosophila Homologues of Adenomatous Polyposis Coli (APC) and the Formin Diaphanous Collaborate by a Conserved Mechanism to Stimulate Actin Filament Assembly. J. Biol. Chem. 288, 13897–13905.

Kang, H., Wang, J., Longley, S.J., Tang, J.X., and Shaw, S.K. (2010). Biochemical and Biophysical Research Communications Relative actin nucleation promotion efficiency by WASP and WAVE proteins in endothelial cells. Biochem. Biophys. Res. Commun. 400, 661–666.

Kelber, J.A., and Klemke, R.L. (2011). The Actin Cytoskeleton.

Kim, B.N., Ahn, D.H., Kang, N., Yeo, C.D., Kim, Y.K., Lee, K.Y., Kim, T.J., Lee, S.H., Park, M.S., Yim, H.W., et al. (2020). TGF-β induced EMT and stemness characteristics are associated with epigenetic regulation in lung cancer. Sci. Rep. 10, 1–11.

Klein, T.J., Jenny, A., Djiane, A., and Mlodzik, M. (2006). CKIε/discs overgrown Promotes Both Wnt-Fz/β-Catenin and Fz/PCP Signaling in Drosophila. Curr. Biol. 16, 1337–1343.

Ko, K.K., Powell, M.S., and Hogarth, P.M. (2014). ZSWIM1□: A novel biomarker in T helper cell differentiation. Immunol. Lett. 160, 133–138.

Koch, N., Dharmalingam, E., Westermann, M., Qualmann, B., Thomas, U., and Kessels, M.M. (2012). Abp1 utilizes the Arp2 / 3 complex activator Scar / WAVE in bristle development. J. Cell Sci. 125, 3578–3589.

Koestler, S.A., Rottner, K., Lai, F., Block, J., Vinzenz, M., and Small, J.V. (2009). F-and G-actin concentrations in lamellipodia of moving cells. PLoS One 4, 3–7.

Korobova, F., and Svitkina, T. (2008). Arp2/3 complex is important for filopodia formation, growth cone motility, and neuritogenesis in neuronal cells. Mol. Biol. Cell 19, 1561–1574.

Kovar, D.R. (2006). Molecular details of formin-mediated actin assembly. Curr. Opin. Cell Biol. 18, 11–17.

Kullmann, J.A., Meyer, S., Pipicelli, F., Kyrousi, C., Schneider, F., Bartels, N., Cappello, S., and Rust, M.B. (2020). Profilin1-Dependent F-Actin Assembly Controls Division of Apical Radial Glia and Neocortex Development. Cereb. Cortex 30, 3467–3482.

Kunda, P., Craig, G., Dominguez, V., and Baum, B. (2003). Abi, Sra1, and Kette Control the Stability and Localization of SCAR / WAVE to Regulate the Formation of Actin-Based Protrusions. Curr. Biol. 13, 1867–1875.

Lee, H., and Adler, P.N. (2004). The grainy head transcription factor is essential for the function of the frizzled pathway in the Drosophila wing. Mech. Dev. 121, 37–49.

Legg, J., Bompard, G., Dawson, J., Morris, H., Andrew, N., Cooper, L., Johnston, S., Tramountains, G., and Machesky, L. (2007). N-WASP involvement in dorsal ruffle formation in mouse embryonic fibroblasts. Mol. Biol. Cell 18, 986–994.

Leung, T., Chen, X.-Q., Tan, I., Manser, E., and Lim, L. (1998). Myotonic Dystrophy Kinase-Related Cdc42-Binding Kinase Acts as a Cdc42 Effector in Promoting Cytoskeletal Reorganization. Mol. Cell. Biol. 18, 130–140.

Liao, M., and Peng, L. (2020). MiR-206 may suppress non-small lung cancer metastasis by targeting CORO1C. Cell. Mol. Biol. Lett. 25.

Lu, Q., and Adler, P.N. (2015). The diaphanous Gene of Drosophila Interacts Antagonistically with multiple wing hairs and Plays a Key Role in Wing Hair Morphogenesis. PLoS One 10, 1–20.

Lu, Q., Yan, J., and Adler, P.N. (2010). The Drosophila planar polarity proteins inturned and multiple wing hairs interact physically and function together. Genetics 185, 549–558.

Lu, Q., Schafer, D.A., and Adler, P.N. (2015). The Drosophila planar polarity gene multiple wing hairs directly regulates the actin cytoskeleton. Development 142, 2478–2486.

Luo, L., Lee, T., Tsai, L., Tang, G., Jan, L.Y., and Jan, Y.N. (1997). Genghis Khan (Gek) as a putative effector for Drosophila Cdc42 and regulator of actin polymerization. Proc. Natl. Acad. Sci. U. S. A. 94, 12963–12968.

Machesky, L.M., Mullins, R.D., Higgs, H.N., Kaiser, D.A., Blanchoin, L., May, R.C., Hall, M.E., and Pollard, T.D. (1999). Scar, a WASp-related protein, activates nucleation of actin filaments by the Arp2/3 complex. Proc. Natl. Acad. Sci. U. S. A. 96, 3739–3744.

Matis, M., Russler-Germain, D.A., Hu, Q., Tomlin, C.J., and Axelrod, J.D. (2014). Microtubules provide directional information for core PCP function. Elife 3, e02893.

Matusek, T., Djiane, A., Jankovics, F., Brunner, D., Mlodzik, M., and Mihály, J. (2006). The Drosophila formin DAAM regulates the tracheal cuticle pattern through organizing the actin cytoskeleton. Development 133, 957–966.

Matusek, T., Gombos, R., Szécsényi, A., Sánchez-Soriano, N., Czibula, Á., Pataki, C., Gedai, A., Prokop, A., Raskó, I., and Mihály, J. (2008). Formin proteins of the DAAM subfamily play a role during axon growth. J. Neurosci. 28, 13310–13319.

Miki, H., Suetsugu, S., and Takenawa, T. (1998). WAVE, a novel WASP-family protein involved in actin reorganization induced by Rac. EMBO J. 17, 6932–6941.

Mitchell, H., Roach, J., and Petersen, N. (1983). The Morphogenesis of Cell Hairs on Drosophila Wings. Dev. Biol. 95, 387–398.

Moreira, C.G.A., Jacinto, A., and Prag, S. (2013). Drosophila integrin adhesion complexes are essential for hemocyte migration in vivo. Biol Open 2, 795–801.

Mullins, R.D., Heuser, J.A., and Pollard, T.D. (1998). The interaction of Arp2/3 complex with actin: Nucleation, high affinity pointed end capping, and formation of branching networks of filaments. Proc. Natl. Acad. Sci. U. S. A. 95, 6181–6186.

Okumura, F., Oki, N., Fujiki, Y., Ikuta, R., Osaki, K., Hamada, S., Nakatsukasa, K., Hisamoto, N., Hara, T., and Kamura, T. (2021). ZSWIM8 is a myogenic protein that partly prevents C2C12 differentiation. Sci. Rep. 11.

de Oliveira Rodrigues, R., Yaochite, J.N.U., Sasahara, G.L., Albuquerque, A.A., da Cruz Fonseca, S.G., de Vasconcelos Araújo, T.D., Santiago, G.M.P., de Sousa, L.M., de Carvalho, J.L., Alves, A.P.N.N., et al. (2020). Antioxidant, anti-inflammatory and healing potential of ethyl acetate fraction of Bauhinia ungulata L. (Fabaceae) on in vitro and in vivo wound model. Mol. Biol. Rep. 47, 2845–2859.

Paunola, E., Mattila, P.K., and Lappalainen, P. (2002). WH2 domain□: a small, versatile adapter for actin monomers. FEBS Lett. 513, 92–97.

Piao, Y.L., Wu, Y., Seo, S.Y., Lim, S.C., and Cho, H. (2014). Wound healing effects of new 15-hydroxyprostaglandin dehydrogenase inhibitors. Prostaglandins Leukot. Essent. Fat. Acids 91, 325–332.

Pollitt, A.Y., and Insall, R.H. (2009a). WASP and SCAR / WAVE proteins□: the drivers of actin assembly. J. Cell Sci. 122, 2575–2578.

Pollitt, A.Y., and Insall, R.H. (2009b). Loss of Dictyostelium HSPC300 causes a scar-like phenotype and loss of SCAR protein. BMC Cell Biol. 10, 1–7.

Rajan, A., Tien, A., Haueter, C.M., Schulze, K.L., and Bellen, H.J. (2009). The Arp2 / 3 complex and WASp are required for apical trafficking of Delta into microvilli during cell fate specification of sensory organ precursors. Nat. Publ. Gr. 11, 815–824.

Ren, N., Zhu, C., Lee, H., and Adler, P.N. (2005). Gene expression during Drosophila wing morphogenesis and differentiation. Genetics 171, 625–638.

Ren, N., He, B., Stone, D., Kirakodu, S., and Adler, P.N. (2006a). The shavenoid gene of Drosophila encodes a novel actin cytoskeleton interacting protein that promotes wing hair morphogenesis. Genetics 172, 1643–1653.

Ren, N., He, B., Stone, D., Kirakodu, S., and Adler, P.N. (2006b). The shavenoid Gene of Drosophila Encodes a Novel Actin Cytoskeleton Interacting Protein That Promotes Wing Hair Morphogenesis. Genetics 172, 1643–1653.

Ren, N., Charlton, J., and Adler, P.N. (2007). The flare Gene, Which Encodes the AIP1 Protein of Drosophila, Functions to Regulate F-Actin Disassembly in Pupal Epidermal Cells. Genetics 176, 2223–2234.

Revenu, C., Athman, R., Robine, S., and Louvard, D. (2004). The co-workers of actin filaments□: from cell structures to signals. Nat. Rev. Mol. Cell Biol. 5, 1–12.

Rodriguez-Mesa, E., Abreu-Blanco, M.T., Rosales-Nieves, A.E., and Parkhurst, S.M. (2012). Developmental expression of Drosophila Wiskott-Aldrich Syndrome family proteins. Dev. Dyn. 241, 608–626.

Rottner, K., Faix, J., Bogdan, S., Linder, S., and Kerkhoff, E. (2017). Actin assembly mechanisms at a glance. J. Cell Sci. 130, 3427–3435.

Rotty, J.D., Wu, C., and Bear, J.E. (2012). New insights into the regulation and cellular functions of the ARP2/3 complex. Nat. Rev. Mol. Cell Biol. 14, 7–12.

Rust, M.B., Kullmann, J.A., and Witke, W. (2012). Role of the actin-binding protein profilin1 in radial migration and glial cell adhesion of granule neurons in the cerebellum. Cell Adhes. Migr. 6, 13–17.

Sagner, A., Merkel, M., Aigouy, B., Gaebel, J., Brankatschk, M., Julicher, F., and Eaton, S. (2012). Establishment of Global Patterns of Planar Polarity during Growth of the Drosophila Wing Epithelium. Curr. Biol. 22, 1296–1301.

Sánchez-Gutiérrez, D., Sáez, A., Pascual, A., and Escudero, L.M. (2013). Topological progression in proliferating epithelia is driven by a unique variation in polygon distribution. PLoS One 8, e79227 1–8.

Santos, J., Silva, D., and Dotti, C.G. (2002). BREAKING THE NEURONAL SPHERE: REGULATION OF THE ACTIN CYTOSKELETON IN NEURITOGENESIS. Nat. Rev. Neurosci. 3, 694–704.

Schneider, I. (1972). Cell lines derived from late embryonic stages of Drosophila melanogaster. J. Embryol. Exp. Morphol. 27, 353–365.

Shellard, A., and Mayor, R. (2020). All Roads Lead to Directional Cell Migration. Trends Cell Biol. 30, 852–868.

Shi, C.Y., Kingston, E.R., Kleaveland, B., Lin, D.H., Stubna, M.W., and Bartel, D.P. (2020). The ZSWIM8 ubiquitin ligase mediates target-directed microRNA degradation. Science (80-.). 370, eabc9359 1–10.

Singh, J., and Mlodzik, M. (2012). Planar Cell Polarity Signaling: Coordination of cellular orientation across tissues. Wiley Interdiscip Rev Dev Biol 1, 479–499.

Stephan, R., Gohl, C., Fleige, A., Klämbt, C., and Bogdan, S. (2011). Membrane-targeted WAVE mediates photoreceptor axon targeting in the absence of the WAVE complex in Drosophila. Mol. Biol. Cell 22, 4079–4092.

Strutt, H., and Strutt, D. (2009). Asymmetric localisation of planar polarity proteins: Mechanisms and consequences. Semin. Cell Dev. Biol. 20, 957–963.

Suarez, C., and Kovar, D.R. (2016). Internetwork competition for monomers governs actin cytoskeleton organization. Nat. Rev. Mol. Cell Biol. 17, 799–810.

Suarez, C., Carroll, R.T., Burke, T.A., Christensen, J.R., Bestul, A.J., Sees, J.A., James, M.L., Sirotkin, V., and Kovar, D.R. (2015). Article Profilin Regulates F-Actin Network Homeostasis by Favoring Formin over Arp2 / 3 Complex. Dev. Cell 8, 1–11.

Swaney, K., and Li, R. (2016). Function and regulation of the Arp2/3 complex during cell migration in diverse environments. Curr. Opin. Cell Biol. 42, 63–72.

Tahirovic, S., Hellal, F., Neukirchen, D., Hindges, R., Garvalov, B.K., Flynn, K.C., Stradal, T.E., Chrostek-grashoff, A., Brakebusch, C., and Bradke, F. (2010). Rac1 Regulates Neuronal Polarization through the WAVE Complex. J. Neurosci. 30, 6930–6943.

Takenawa, T., and Suetsugu, S. (2007). The WASP–WAVE protein network: connecting the membrane to the cytoskeleton. Nat. Rev. Mol. Cell Biol. 8, 37–48.

Tirouvanziam, R., Davidson, C.J., Lipsick, J.S., and Herzenberg, L.A. (2004). Fluorescence-activated sorting (FACS) of Drosophila hemocytes reveals important functional similarities to mammalian leukocytes. Proc. Natl. Acad. Sci. U. S. A. 101, 2912–2917.

Turner, C.M., and Adler, P.N. (1998). Distinct roles for the actin and microtubule cytoskeletons in the morphogenesis of epidermal hairs during wing development in Drosophila. Mech. Dev. 70, 181–192.

Vignjevic, D., Kojima, S., Aratyn, Y., Danciu, O., Svitkina, T., and Borisy, G.G. (2006). Role of fascin in fi lopodial protrusion. J. Cell Biol. 174, 863–875.

Vitriol, E.A., McMillen, L.M., Kapustina, M., Gomez, S.M., Vavylonis, D., and Zheng, J.Q. (2015). Two functionally distinct sources of actin monomers supply the leading edge of lamellipodia. Cell Rep. 11, 433–445.

Wang, Y., Wang, H., Pan, T., Li, L., Li, J., and Yang, H. (2017). STIM1 silencing inhibits the migration and invasion of A549 cells. Mol. Med. Rep. 16, 3283–3289.

Wang, Z., Hou, Y., Guo, X., van derVoet, M., Boxem, M., Dixon, J.E., Chisholm, A., and Jin, Y. (2013). The EBAX-type Cullin-RING E3 Ligase and Hsp90 Guard the Protein Quality of the SAX-3/Robo receptor in developing neurons. Neuron 79, 903–916.

Watanabe, N., Madaule, P., Reid, T., Ishizaki, T., Watanabe, G., Kakizuka, A., Saito, Y., Nakao, K., Jockusch, B.M., and Narumiya, S. (1997). p140mDia, a mammalian homolog of Drosophila diaphanous, is a target protein for Rho small GTPase and is a ligand for profilin. EMBO J. 16, 3044–3056.

Winter, C.G., Wang, B., Ballew, A., Royou, A., Karess, R., Axelrod, J.D., and Luo, L. (2001). Drosophila Rho-associated kinase (Drok) links Frizzled-mediated planar cell polarity signaling to the actin cytoskeleton. Cell 105, 81–91.

Wong, L.L., and Adler, P. (1993). Tissue Polarity Genes of Drosophila regulate the subcellular location for prehair initiation in pupal wing cells. J. Cell Biol. 123, 209–221.

Wu, J.S., and Luo, L. (2006). A protocol for mosaic analysis with a repressible cell marker (MARCM) in Drosophila. Nat. Protoc. 1, 2583–2590.

Yamamoto, S., Jaiswal, M., Charng, W.L., Gambin, T., Karaca, E., Mirzaa, G., Wiszniewski, W., Sandoval, H., Haelterman, N.A., Xiong, B., et al. (2014). A drosophila genetic resource of mutants to study mechanisms underlying human genetic diseases. Cell 159, 200–214.

Yan, J., Huen, D., Morely, T., Johnson, G., Gubb, D., Roote, J., and Adler, P.N. (2008). The multiple-wing-hairs gene encodes a novel GBD-FH3 domain-containing protein that functions both prior to and after wing hair initiation. Genetics 180, 219–228.

Yan, J., Lu, Q., Fang, X., and Adler, P.N. (2009). Rho1 has multiple functions in Drosophila wing planar polarity. Dev. Biol. 333, 186–199.

Yaniv, S.P., Meltzer, H., Alyagor, I., and Schuldiner, O. (2020). Developmental axon regrowth and primary neuron sprouting utilize distinct actin elongation factors. J. Cell Biol. 219.

Zallen, J.A., Cohen, Y., Hudson, A.M., Cooley, L., Wieschaus, E., and Schejter, E.D. (2002). SCAR is a primary regulator of Arp2/3-dependent morphological events in Drosophila. J. Cell Biol. 156, 689–701.

Zhao, K., Wang, D., Zhao, X., Wang, C., Gao, Y., Liu, K., Wang, F., Wu, X., Wang, X., Sun, L., et al. (2020). WDR63 inhibits Arp2/3-dependent actin polymerization and mediates the function of p53 in suppressing metastasis. EMBO Rep. 21, 1–13.

