## Supplemental Data for "The conserved Pelado/ZSWIM8 protein regulates actin dynamics by promoting linear actin filament polymerization"

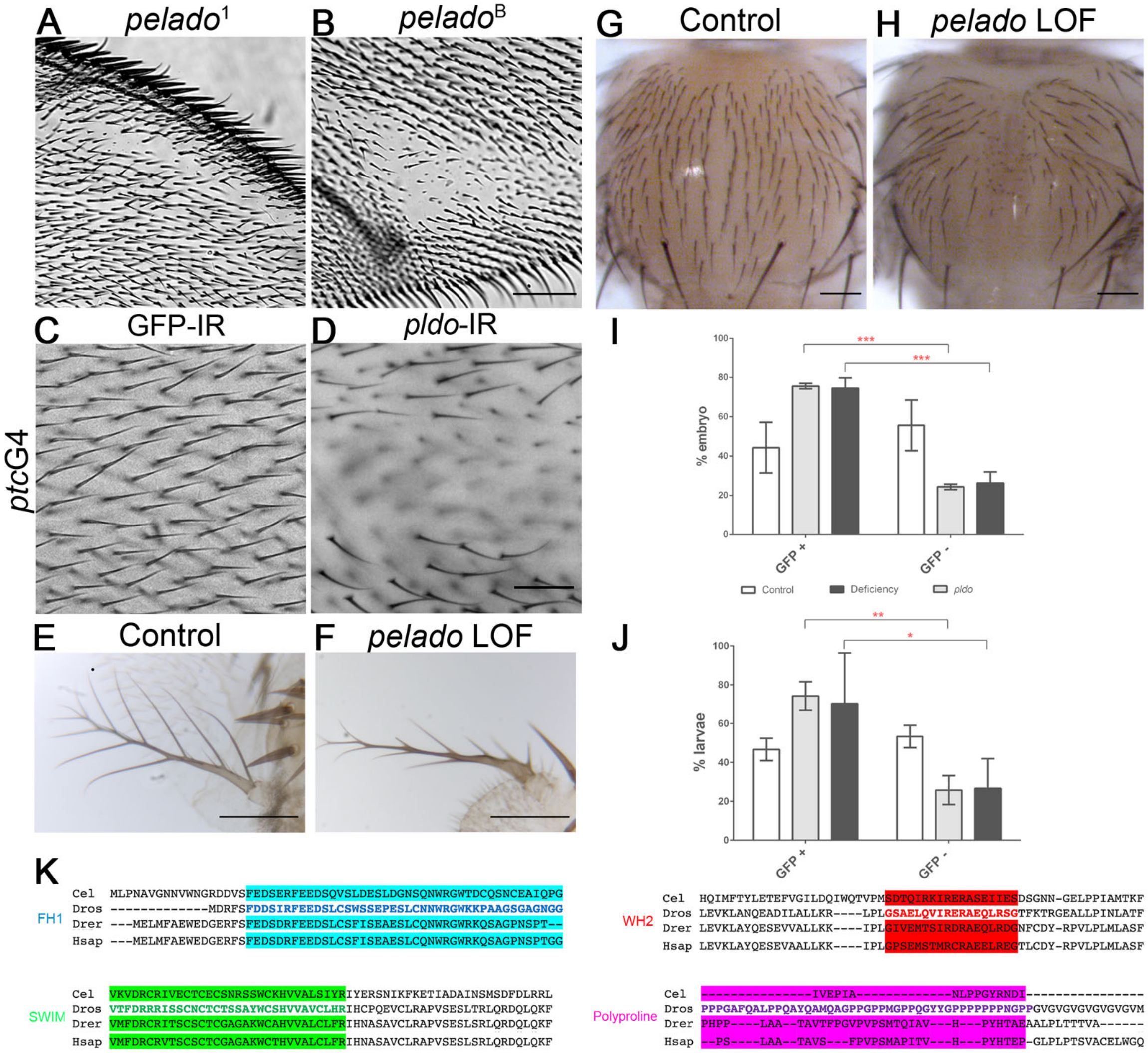


**Suppl Fig S1: Pldo is required for hair and laterals formation.**

A-B. Characterization of CG34401/*pldo* LOF phenotypes. Adult wings with unmarked *pldo* clones showing the absence of hairs phenotype. In these cases, different alleles for *pldo* were used (*pldo*$^1$ and *pldo*$^B$). Scale bar correspond to 100 μm.

C-D. Adult wings showing the *pldo* LOF phenotype generated by the expression of RNAi (and RNAi against GFP as a control) in the *ptcGal4* expression domain in the wing (note that *ptc* is expressed between veins 2 and 3 in the wing along the A/P compartment boundary, see schematic of *ptc* expression in Figure 3). Scale bar correspond to 50 μm.

E-F. *pldo* LOF phenotype in adult antenna, generated by *ey-Gal4* driven RNAi expression (and control RNAi for comparison*)*. Scale bars correspond to 50 μm.

G-H. Adult thorax showing the absence of cuticular hairs generated by the expression of RNAi (and RNAi against GFP as a control) in the *pannierGal4* expression domain in the notum (note that bristles are not affected). Scale bars corresponds to 100 μm.

I-J. Quantification of embryo and larvae mutants for *pldo* (GFP negative) or control (GFP positive). In each experiment, at least 20 embryo/larvae were considered per genotype, n=3. 2-way ANOVA plus Sidak's multiple comparison test was used for statistical analysis, **** indicates P<0,0001; ** indicates P=0,0039 and * indicates P=0,0259.

K. *pldo* is located on the X chromosome at position X: 18,863,950… X: 18,884,075, and BLAST analysis indicates that *pldo* is conserved from *C. elegans* to humans. Protein sequence alignment of main predicted domains from different species: *Cel*: *C. elegans, Dros: Drosophila, Drer: Danio rerio,* and *Hsap*: *Homo sapiens*. Alignment was obtained through the software package Clustal Omega (RRID:001591).

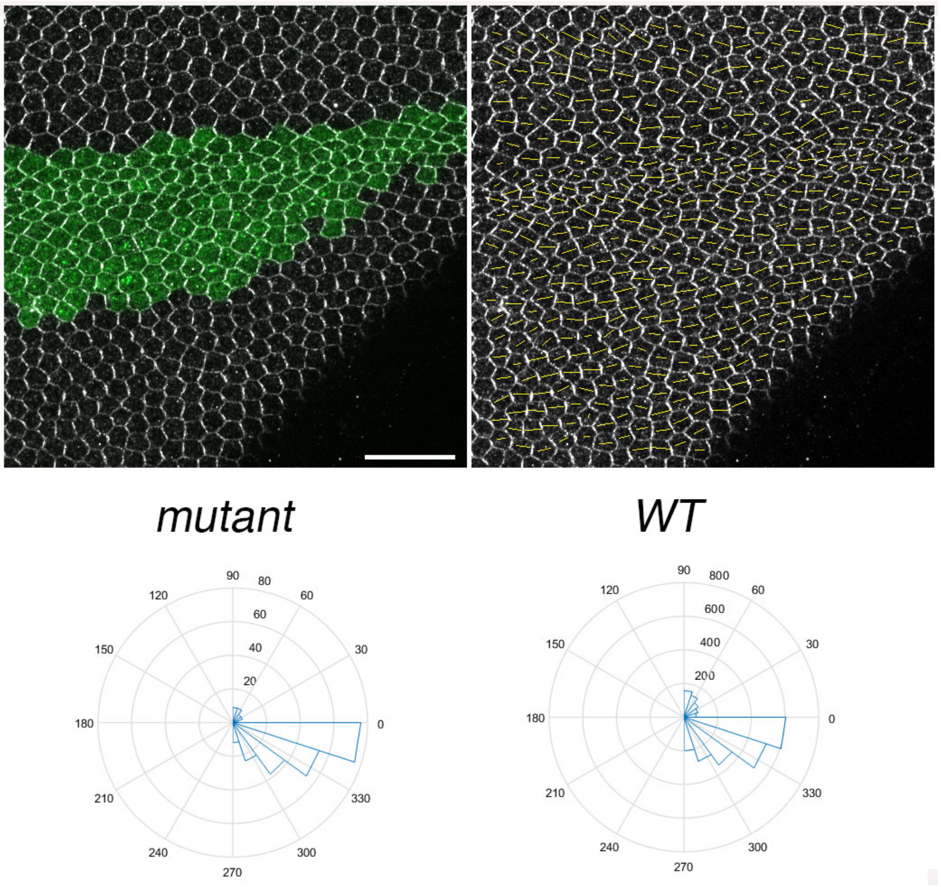


**Suppl Fig S2: Pldo does not affect the core Fz/PCP pathway.**

Asymmetric Flamingo (Fmi) localization, as a proxy for Fz/PCP signaling, is shown. Note that it is not affected in *pldo* mutant cells, as compared to surrounding control *wt* cells, in pupal wings (as shown in the graphs). Clones of mutant cells were generated by the MARCM technique, with mutant cells marked with GFP (green). Polarity strength lines of Fmi staining is shown as orange lines. 6 different individuals were used, and around 100 cells were analyzed in each case. Scale bar represents 100 μm.

A

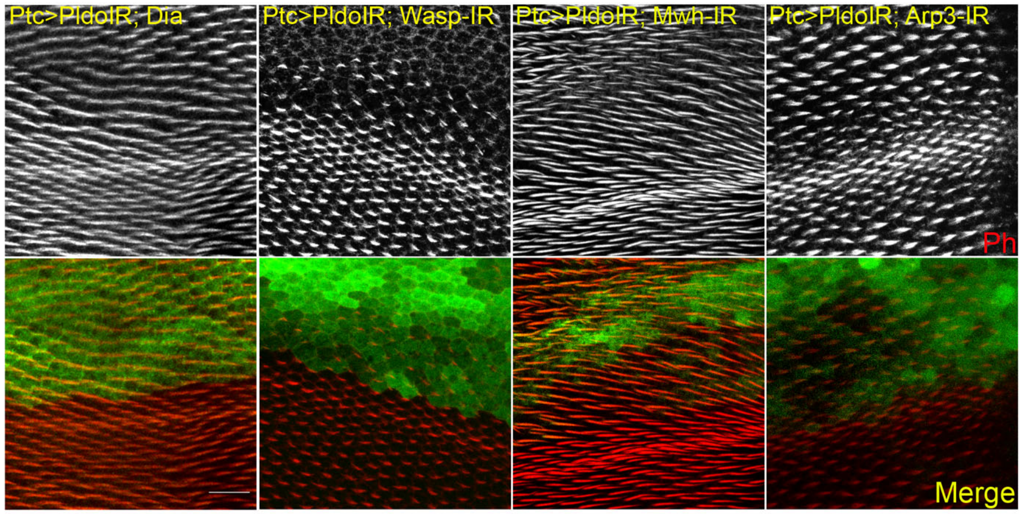


B

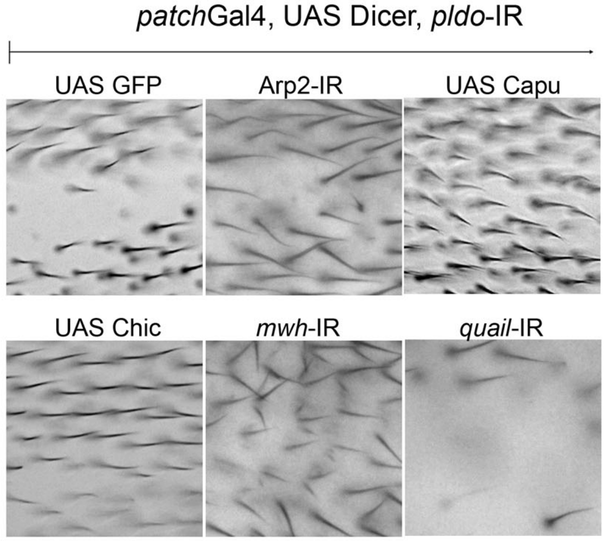


C

| Genotype | Rescue |
|---|---|
| PtcG4, Dcr, PldoIR x GFP (IR) | no |
| PtcG4, Dcr, PldoIR x Act | yes |
| PtcG4, Dcr, PldoIR x Dia | yes |
| PtcG4, Dcr, PldoIR x Scar-IR | yes |
| PtcG4, Dcr, PldoIR x Scar-GFP | no |
| PtcG4, Dcr, PldoIR x Scar-Myr | no |
| PtcG4, Dcr, PldoIR x Mwh-IR | yes |
| PtcG4, Dcr, PldoIR x Arp2-IR | yes |
| PtcG4, Dcr, PldoIR x Wasp-IR | no |
| PtcG4, Dcr, PldoIR x Arp3-IR | yes |
| PtcG4, Dcr, PldoIR x Abi | partial |
| PtcG4, Dcr, PldoIR x Ena | yes |
| PtcG4, Dcr, PldoIR x Sha | yes |
| PtcG4, Dcr, PldoIR x Capu | no |
| PtcG4, Dcr, PldoIR x FL-Daam | no |
| PtcG4, Dcr, PldoIR x N-Daam | no |
| PtcG4, Dcr, PldoIR x C-Daam | no |
| PtcG4, Dcr, PldoIR x Daam-IR | no |
| PtcG4, Dcr, PldoIR x CA-tsr | no |
| PtcG4, Dcr, PldoIR x I-tsr | no |
| PtcG4, Dcr, PldoIR x Tsr | no |

**Suppl Fig S3: Pldo mediates competition for monomeric Actin during trichome formation, extended.**

A. *pldo* LOF phenotype reversion evaluated in pupal and adult wings. Confocal images of pupal wings expressing *ptcGal4*, *UAS-Dcr*, *UAS-pldo*IR in combination with the indicated transgenes. Rhodamine conjugated to phalloidin was used to stain actin filaments and GFP was used to mark the *ptcGAL4* domain in the wing. Scale bar correspond to 50 μm.

B. Additional adult wings showing the interaction between *ptcGal4*, *UAS-Dcr*, *UAS-pldo*IR and the indicated transgenes (see Fig. 3 C for quantification). Quantification was performed as described in main text Figure 3C. Scale bar correspond to 50 μm.

C. Table displaying all interactions assayed with *ptcGAL4, UAS-Dcr, UAS-pldo*IR. Data are based on scoring at least 10 individuals in each case.

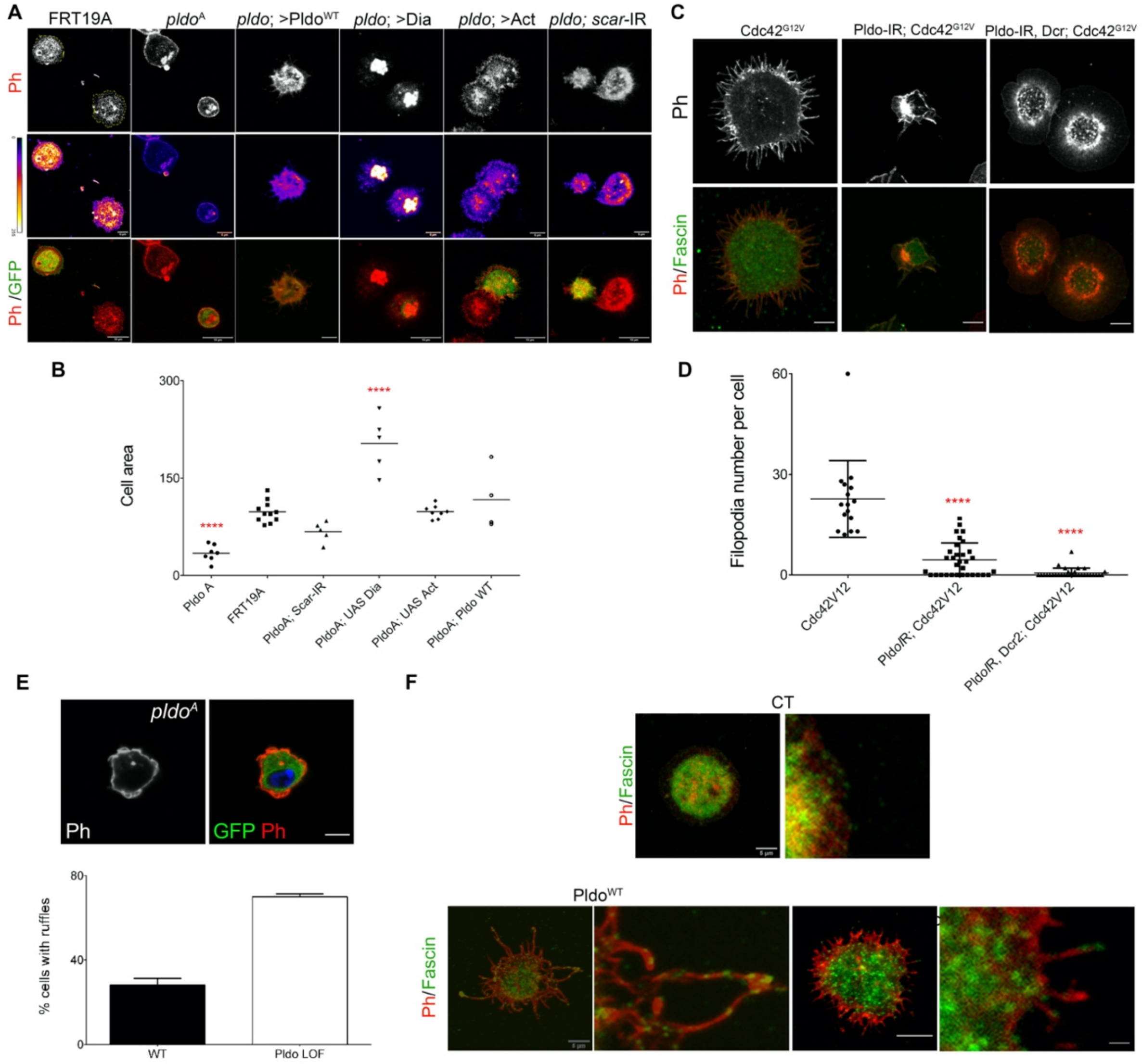


**Suppl Fig S4: Pldo is essential to induce filopodia formation and to sustain cell shape in hemocytes.**

A. *pldo* mutant cells were generated by the MARCM technique (expressing GFP, green cells). An FRT19A chromosome was used as a control. Note that *pldo* mutant cells show a reduction in the cell attachment area, that was rescued by the expression of either Pldo, Dia or Act, or by knocking-down *scar*. Scale bars correspond to 10 µm.

B. Cell area was quantified using the Phalloidin staining (red and heatmap), using at least 10 cells per genotype in each experiment (and 3 independent experiments). Statistical analysis was 1-Way ANOVA, Tukey post-test *P*<0,0001.

C-D. Filopodia formation induced by the expression of $Cdc42^{G12V}$ in hemocytes, was evaluated by knocking down *pldo*. With or without Dicer expression in the LOF of Pldo, there was a significant reduction in filopodia formation by $Cdc42^{G12V}$. Scale bars represents 5 µm. Number of filopodia per cell was quantified in at least 10 cells per genotype in each experiment (and 3 independent experiments). Statistical analysis was 1-Way ANOVA, Tukey post-test $P$<0,0001.

E. Ruffle formation in *pldo* mutant cells. Ruffle presence was quantified in at least 10 cells per genotype in each experiment (3 indep. experiments). Scale bar represents 5 µm.

F. Fascin (green, staining bundled linear actin) serves as a marker for mature filopodia. Higher magnification of filopodia showing the Fascin presence in the distal filopodial region in Pldo GOF or Cdc42G12V. Scale bar represents 5µm and 20µm in high magnification views.

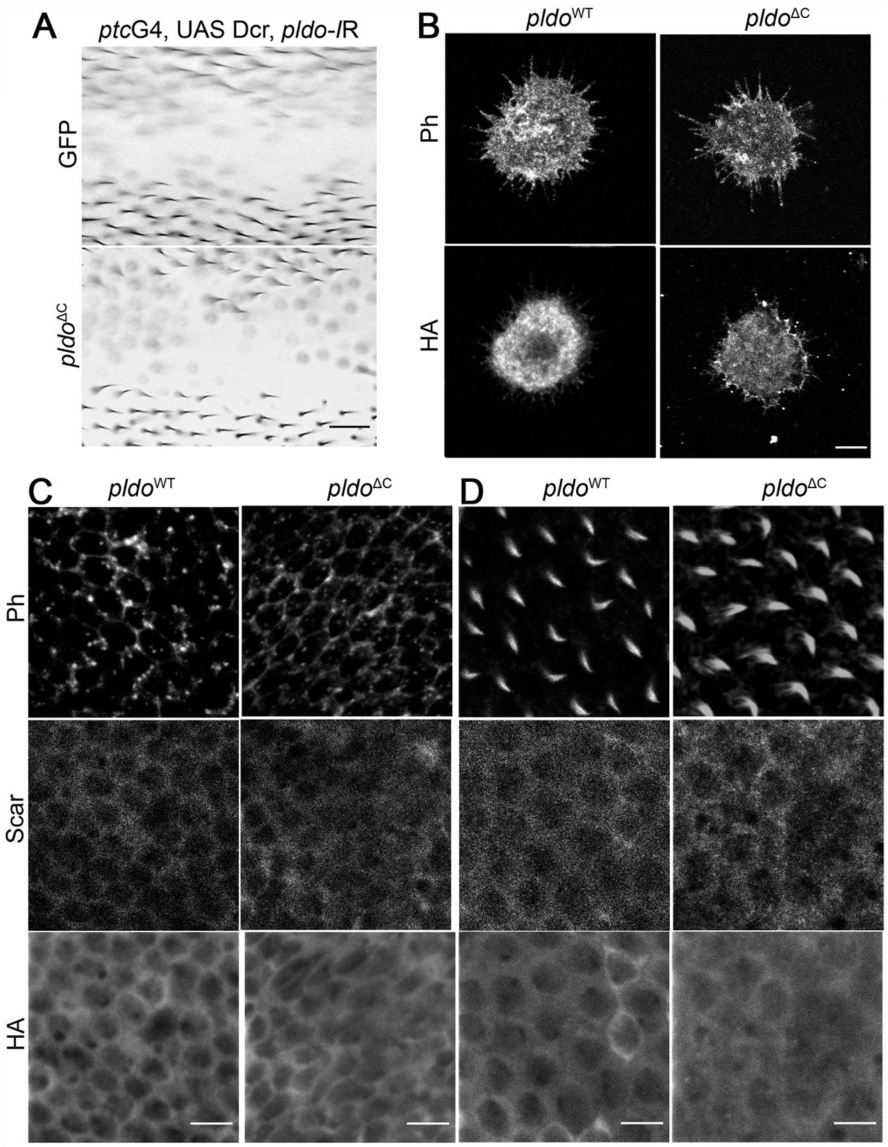


**Suppl Fig S5: N-terminal region of Pldo is sufficient to induce filopodia formation.**

A. Pldo$^{\Delta C}$ was not capable to rescue the lack of trichome phenotype observed by *pldo* LOF in adult wings, when co-expressing with the *UAS-pldo*-IR in the *ptcGAL4* expression domain. No change in the phenotype was observed, confirming the *pldo* LOF mutant defects, as seen in pupal wings (Figure 5D). Scale bar represents 50 μm.

B. Localization of Pldo was evaluated in confocal microscopy images, through the HA-tag staining, in hemocytes, observing no notable differences. These experiments were performed at 18ºC, with near wild-type expression levels to prevent overexpression of the transgenes. Scale bar represents 5μm.

C-D. Localization of Pldo was evaluated in confocal microscopy images, through the HA-tag staining, in pupal wings, before and after hair formation, observing no notable differences. These experiments were performed at 18ºC, at 64 h (C) or 72 h (D) APF. With near wild-type expression levels to prevent overexpression of the transgenes. Scale bar represents 25 μm.

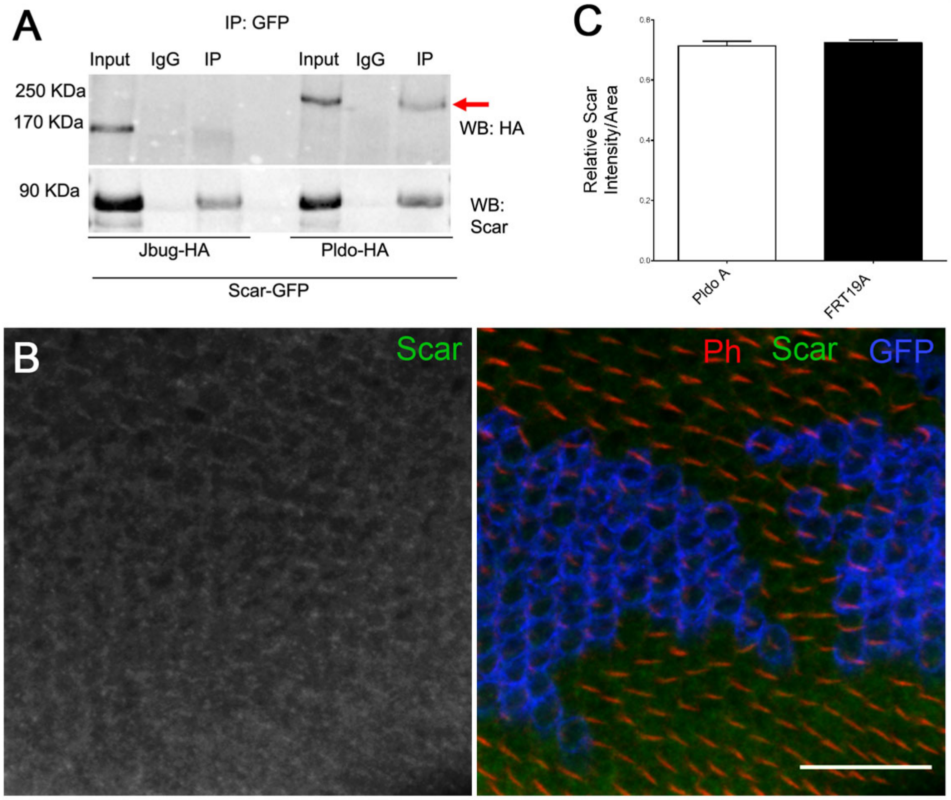


**Suppl Fig S6: Pldo function in actin cytoskeleton regulation is not related to the ubiquitin ligase complex activity.**

A. Two independent publications have shown that the Pldo homologues EBAX-1/ZSWIM8 form part of a ubiquitin ligase complex serving an E3 ligase function, recognizing the substrate targets to be degraded by the proteasome. We did not observe such a role of the gene in the actin dynamics context. IP assay evaluating the interaction of Pldo and Scar. S2 cells were co-transfected with either Pldo-HA and Scar-GFP and were immunoprecipitated with anti-GFP. Jitterbug-HA was used as a control (Jbug-HA). Note Scar immunoprecipitated with $Pldo^{WT}$, suggesting that they form a complex. A complete sample gel of this IP is shown in Suppl Figure S7.

B-C. Scar was tested as a potential target of the ubiquitin ligase complex, but no detectable differences on Scar levels were observed. At least 3 individuals were considered in each experiment and each condition: independent experiments n=3. Statistical analysis was done via t-test. Scale bars correspond to 100 µm. We also evaluated Scar levels using MG132 (proteasome inhibitor) and again, we did not observe any significant change in Scar levels, but we observed (in the same gel) an increase in p53 protein levels when cells were incubated with MG132, indicating that the drug was functional.

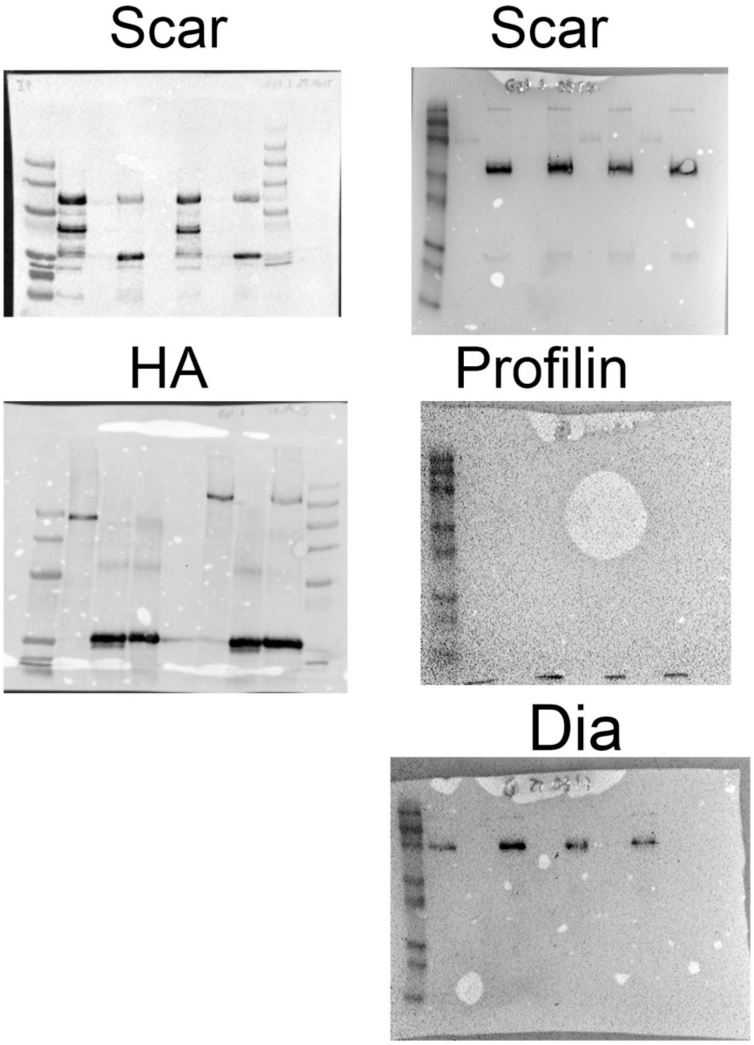


**Suppl Fig S7: IP gels.**

Full size gels from the IP experiments shown above and in main text. On the left are gels that correspond to Suppl Figure S6. On the right are the gels that correspond to Figure 6.
